# Evidence-constrained mechanistic synthesis for pre-calibration perturbation inference

**DOI:** 10.64898/2026.08.17.745376

**Authors:** Dipayan Sengupta, Saumya Panda

**Affiliations:** Consultant Dermatologist, Charnock Hospital, Kolkata, India; Professor and Head, Department of Dermatology, JIMS, Kolkata, India

## Abstract

Biomedical evidence can become mechanistically informative before it becomes sufficiently commensurate to calibrate one system model. We developed evidence-constrained mechanistic synthesis (ECMS), a pre-calibration framework that converts only the mathematical information supported by each finding into constraints on a possibility space of executable mechanistic worlds. We reconstructed historical ECMS systems for alopecia areata (AA) and chronic spontaneous urticaria (CSU) using knowledge available by 31 December 2023, froze molecule/regimen inputs, observation semantics and finite ensembles before outcome reveal, and evaluated later studies at prespecified information-property resolution rather than as calibrated clinical predictions. Among 41 registered atomic properties, ECMS committed to 34 executable propositions: 29 were empirically concordant, one was clearly contradicted, and four could not be fully assessed because intermediate reporting or intervention isolation was insufficient; seven additional properties were prespecified abstentions and were not counted as empirical successes. An evidence-lineage audit further separated nine direct historical analogues from 16 near transfers, four mechanistic transfers and four cross-stream syntheses. Baricitinib withdrawal in AA was a cross-stream synthesis: no pre-cutoff baricitinib-withdrawal result was used, yet the frozen system reproduced early persistence followed by progressive loss. Selective MRGPRX2 antagonism in CSU transferred from pre-cutoff route-separation evidence, whereas patient-level clinical efficacy was withheld. A later baricitinib trajectory contradicted the frozen week-36-to-52 monotonicity property, preserving falsifiability. These results show that heterogeneous pre-calibration evidence can be made executable and externally challenged without being converted into unsupported coefficients, posterior biological probabilities or patient-response models.

## Main

Target and route decisions are often required before a programme has the matched exposure, biomarker and clinical-response data needed to fit a system-level model. Human genetic support is associated with higher probabilities of clinical success, illustrating the value of stronger causal evidence for target selection, but most disease programmes must also reason from interventions, longitudinal cohorts, tissue perturbations, ex vivo experiments, comparative pharmacology and molecular studies that constrain different quantities.^1,2^

Mechanistic systems modelling and quantitative systems pharmacology (QSP) provide a powerful downstream regime when relevant components can be calibrated or qualified for a declared context of use. The earlier regime is different: one study may identify topology, another a direction or ordering, another a time boundary, and an observational association may constrain measurement semantics without identifying causality. Treating these heterogeneous findings as measurements of one common coefficient manufactures precision; ignoring them until full calibration discards real information.^3,4^

ECMS addresses this gap through minimum commitment. Each atomic finding is translated only to the strongest mathematical statement it directly supports. The reason is information-theoretic: a stronger statement removes more possibilities. In a toy 100-state space, A>B retains 45 states (1.15 bits), A-B>=3 retains 28 (1.84 bits), whereas an exact state retains one (6.64 bits). The more restrictive statement is not intrinsically better; it demands correspondingly stronger evidence.^5,6,7,8^

The resulting scientific object is a possibility space H(E), not one fitted graph. Biological labels are unresolved sets, and a coarse relation is represented by a bidirectional possibility boundary A <-> X_AB <-> B. X_AB denotes admissible latent completions rather than one hidden node. The boundary exposes possible interfaces in both directions; each complete world instantiates an evidence-admissible directed realization, function family, intensity, context and temporal/history behaviour. Thus the same declared boundary can contain a forward path, reverse path, shared latent cause, feedback or partially resolved mechanism without asserting all of them simultaneously. Full definitions, including non-Markovian history semantics and hierarchical world generation, are given in Supplementary Methodology (Supplementary Note 1).

Here we test whether this discipline can be challenged in a retrospective outcome-blind historical reconstruction. We reconstructed AA and CSU from knowledge available by 31 December 2023, froze evidence allocation and executable ensembles, registered post-cutoff perturbation queries before outcome scoring, traced every later proposition back to its pre-cutoff evidence ancestry, and then asked whether later experiments supported, delimited or falsified the information that had actually become executable.

### ECMS converts heterogeneous findings into an executable possibility space

An ECMS evidence object records provenance, biological context, manipulation or exposure, readout, dependency, transportability and the highest information level the source can support. Hard fences remove worlds that violate non-negotiable scientific constraints. Softer evidence acts through q, an evidence-derived allocation-frequency target rather than a biological probability: an event may be represented in, for example, 0.7 of the evidence-adjusted ensemble without calling 0.7 the probability that the event is biologically true. The least-committal reallocation is an information projection relative to a declared reference measure; its derivation and conditional-margin treatment are in Supplementary Methodology Sections 2-6.5,6,7,8

World generation is structural first. Context and active membership of unresolved sets are sampled before boundary realization; latent subgraph and overlap choices precede function and time/history families; numerical gains, thresholds and input/observation maps are drawn only after the relevant structure exists. This hierarchical Monte Carlo construction prevents a numerical prior from silently deciding whether a relation exists. Each finished world is then executable and audited against all hard fences and evidence-allocation constraints.

The decision layer is separate. Researchers can later specify a desired biological profile and compare package-declared perturbations on matched worlds, but changing that objective cannot change H(E), q or ensemble membership. This separation is important because preference is not evidence and because objective-fit is not intrinsic drug potency or clinical efficacy.

**Figure 1.**
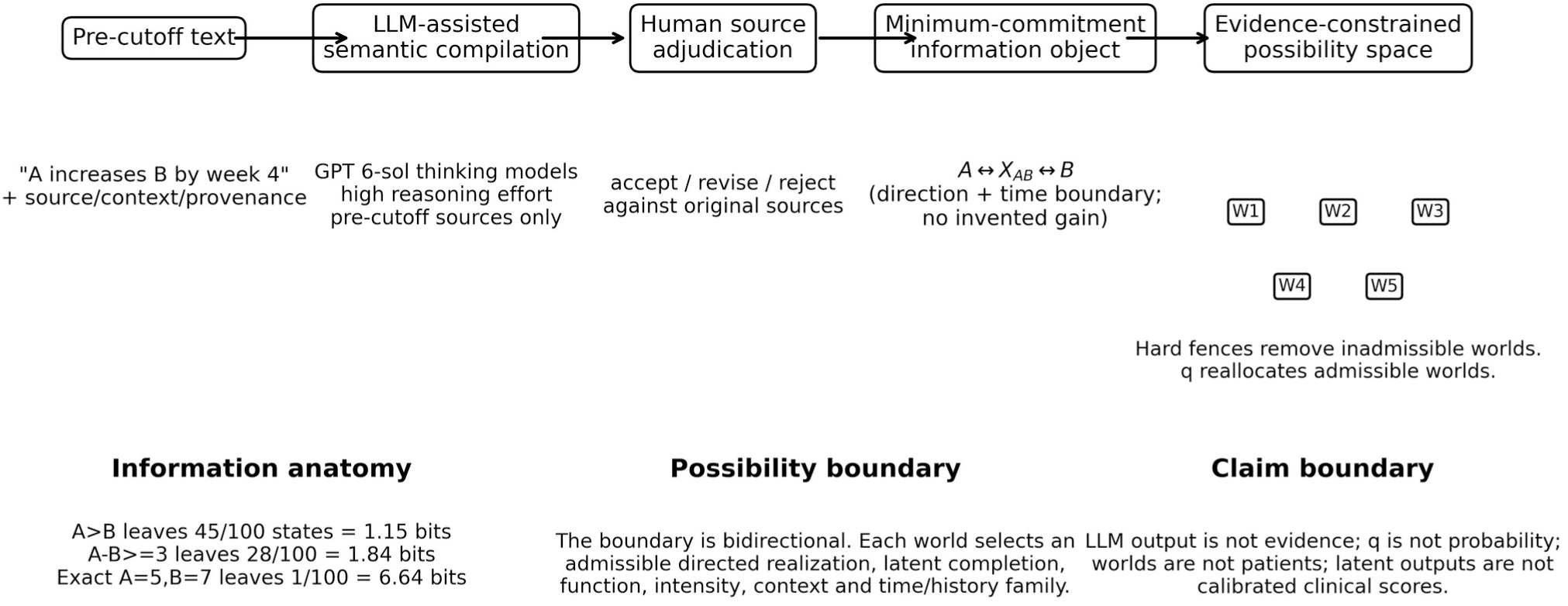
From pre-cutoff language to an evidence-constrained ensemble. A pre-cutoff statement is semantically compiled by an LLM-assisted workflow, manually adjudicated against the original source, translated to the minimum-commitment information object and allocated across admissible whole mechanistic worlds. The A <-> X_AB <-> B boundary is bidirectional while each world realizes an admissible directed subset. LLM output is not evidence; q is not biological probability; worlds are not patients. Detailed mathematics and governance are in Supplementary Methodology and Supplementary Note 2.

#### Box 1 What ECMS quantities do - and do not – mean

Box 1 | What ECMS quantities do - and do not - mean
| Expression | Reader-facing interpretation |
| --- | --- |
| $q=0.80$ | 80% of applicable epistemic worlds are targeted to carry the event under the declared evidence allocation; this is not an 80% probability that the mechanism is true. |
| 75% of worlds improve | 75% of sampled mechanistic hypotheses show that property; this is not a patient-response rate. |
| $A \leftrightarrow X_{AB} \leftrightarrow B$ | Both directional interfaces are available inside the possibility boundary; a given world activates only an admissible directed realization. |
| $X_{AB}$ | A family of possible latent completions, not one unnamed mediator. |
| Abstention | The requested causal/rank/clinical statement is not identified; abstention is not equality, no effect or a successful empirical prediction. |
| Objective-fit % | A downstream research decision-layer loss score, not potency, efficacy, safety or dose optimization. |

### Two historical disease systems were frozen before outcome reveal

The historical cutoff was 31 December 2023. Crucially, the cutoff applied not only to papers but also to concepts, ontology bridges, biological intuition and external reasoning capable of changing H(E). Earliest public availability, including registries, conference disclosures, e-publication and topline releases, determined whether a result was genuinely unseen (Supplementary Note 2).

Because the reconstruction itself was performed in 2026, semantic compilation and biological-intuition candidate generation were assisted between 1 and 20 August 2026 by OpenAI GPT 6-sol thinking models operated at high reasoning effort in an iterative workflow. The models were restricted to supplied pre-cutoff source material for construction tasks and proposed information-anatomy classes, ontology bridges or executable rules for human adjudication; they were not scientific evidence and did not make final acceptance decisions. Investigators manually checked proposals against the cited sources, accepted/revised/rejected them, and manually verified the resulting frozen packages and manuscript-facing intuition log. This governance improves traceability and reduces retrospective leakage risk, but it does not prove that retrospective investigator bias was impossible (Supplementary Note 2; Supplementary Table S18).

The AA atlas contained 106 primary sources and 260 atomic findings; 49 findings carried explicit contradiction or context flags. The CSU atlas contained 81 primary sources and 127 atomic findings, with 44 such flags. Each disease generated 100,000 candidate mechanistic worlds before evidence allocation and equal-weight freezing. AA froze at N=16,384 and CSU at N=32,768, chosen by q fidelity, hard-fence compliance, effective sample size, multiseed reproducibility and standardized benchmark-output stability rather than by a universal ensemble-size rule (Supplementary Notes 3-5; Supplementary Tables S1, S4-S6).

Validation followed the same firewall in both diseases: historical evidence and construction registries -> evidence allocation -> frozen ensemble -> held-out input/observation query -> cryptographic prediction freeze -> outcome reveal. Molecule/regimen identity, route, time and observation semantics were fixed before downstream scoring. Latent hair output was not calibrated to SALT, and latent CSU activity was not calibrated to UAS7 or UCT; nominal doses of different molecules were not mapped onto a common engagement axis (Supplementary Notes 6-7).

**Figure 2.**
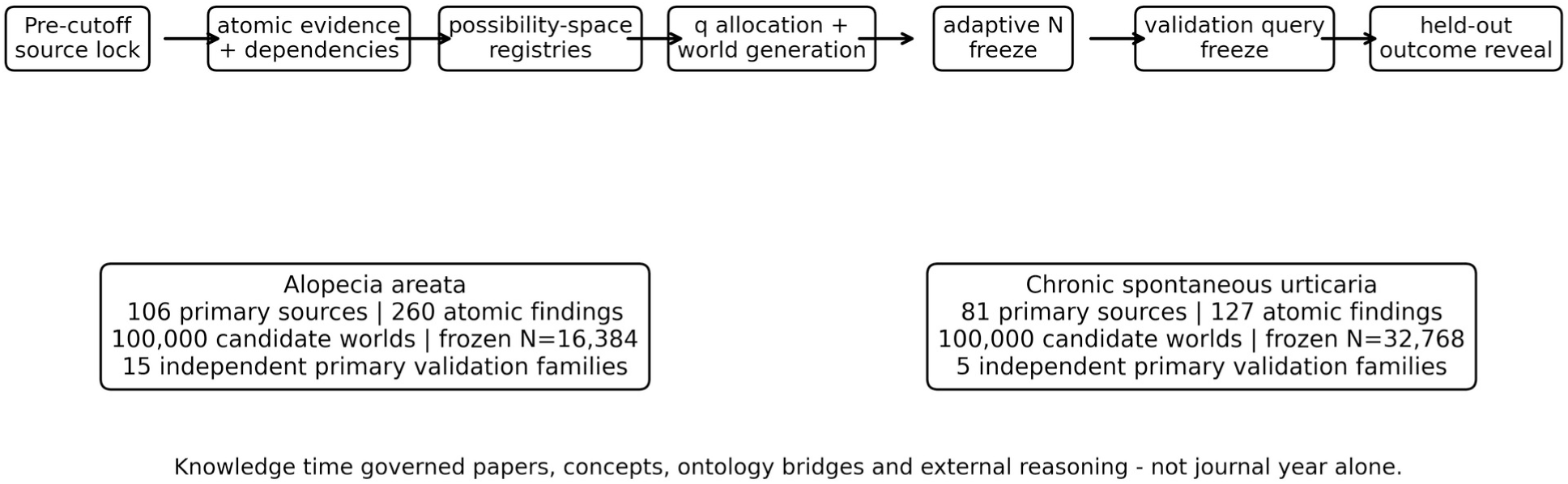
Historical reconstruction and outcome firewall. Two independent disease systems were constructed under the same general ECMS rules and frozen before held-out outcomes were used. The knowledge-time firewall governed scientific concepts and external reasoning as well as source dates. Complete search/adjudication records and freeze identities are provided in Supplementary Notes 2-7 and the reproducibility inventory.

### Evidence ancestry separates replay from mechanistic generalization

The central audit asked a stricter question than whether a later prediction was correct: what pre-cutoff evidence made that proposition executable? Each held-out property was traced backwards through the frozen validation query, input and observation maps, world-level q events and hard fences to pre-cutoff atomic evidence rows, dependency programmes and source identities. We then classified the lineage without assigning a numerical novelty score (Supplementary Note 10; Supplementary Tables S7-S10).

Across 41 atomic properties, ECMS made 34 executable commitments: 29 were empirically concordant, one was a clear mismatch, and four were not fully assessable from the public held-out record because required intermediate values or isolated intervention outcomes were unavailable. The remaining seven were prespecified abstentions and are reported separately rather than counted as successes. Evidence ancestry then divided the same properties into nine direct historical analogues, 16 near transfers, four mechanistic transfers, four cross-stream syntheses, seven abstention boundaries and one mismatch.

These are descriptive lineage classes, not accuracy or novelty scores. The lineage audit also serves as an internal nearest-historical-analogue baseline: direct analogues identify propositions that simple historical matching could plausibly reproduce, whereas transfer and synthesis classes require information beyond exact study/regimen matching. This does not establish superiority to every possible rule-based or fixed-graph baseline; formal matched computational benchmarking remains future work (Supplementary Note 10; Supplementary Tables S9-S11).

Baricitinib withdrawal illustrates cross-stream synthesis. Pre-cutoff evidence established active baricitinib response and continued maturation through week 52, while independent JAK programmes supplied withdrawal/history information. No baricitinib-withdrawal outcome was public before the cutoff. The frozen system nevertheless predicted short-term retention followed by progressive loss after control removal; the later BRAVE-AA1 randomized withdrawal study showed the same qualitative class.^18^ Full lineage: Supplementary Note 10; Supplementary Table S9, row AA-PUB-003.P1.

MRGPRX2 provides a different form of transfer. Pre-cutoff agonist/knockout experiments established an IgE-independent mast-cell route and its separation from the Fc-epsilon-RI route. The held-out interventions were selective antagonists, not repeats of those experiments. Both later antagonist programmes suppressed MRGPRX2-driven degranulation without requiring suppression of the parallel Fc-epsilon-RI route, while the frozen system abstained from converting those mechanistic assays into patient-level CSU efficacy.^21,22^ Full lineage: Supplementary Note 10; Supplementary Table S9, rows CSU-PUB-127.P1/P2 and CSU-PUB-128.P1/P2.

The sole explicit lineage mismatch is equally informative. The frozen AA system encoded continued week-36-to-52 maturation under baricitinib 4 mg because that same-regimen temporal property existed historically. A later real-world cohort improved through week 36 but mean SALT worsened from 37.77 at week 36 to 42.41 at week 52.^19^ Thus the failed atomic proposition was not a speculative extrapolation; it contradicted a directly inherited historical property. Full lineage and grade: Supplementary Note 8 and Supplementary Note 10; Supplementary Tables S7-S9, row AA-PUB-099.P1.

**Figure 3.**
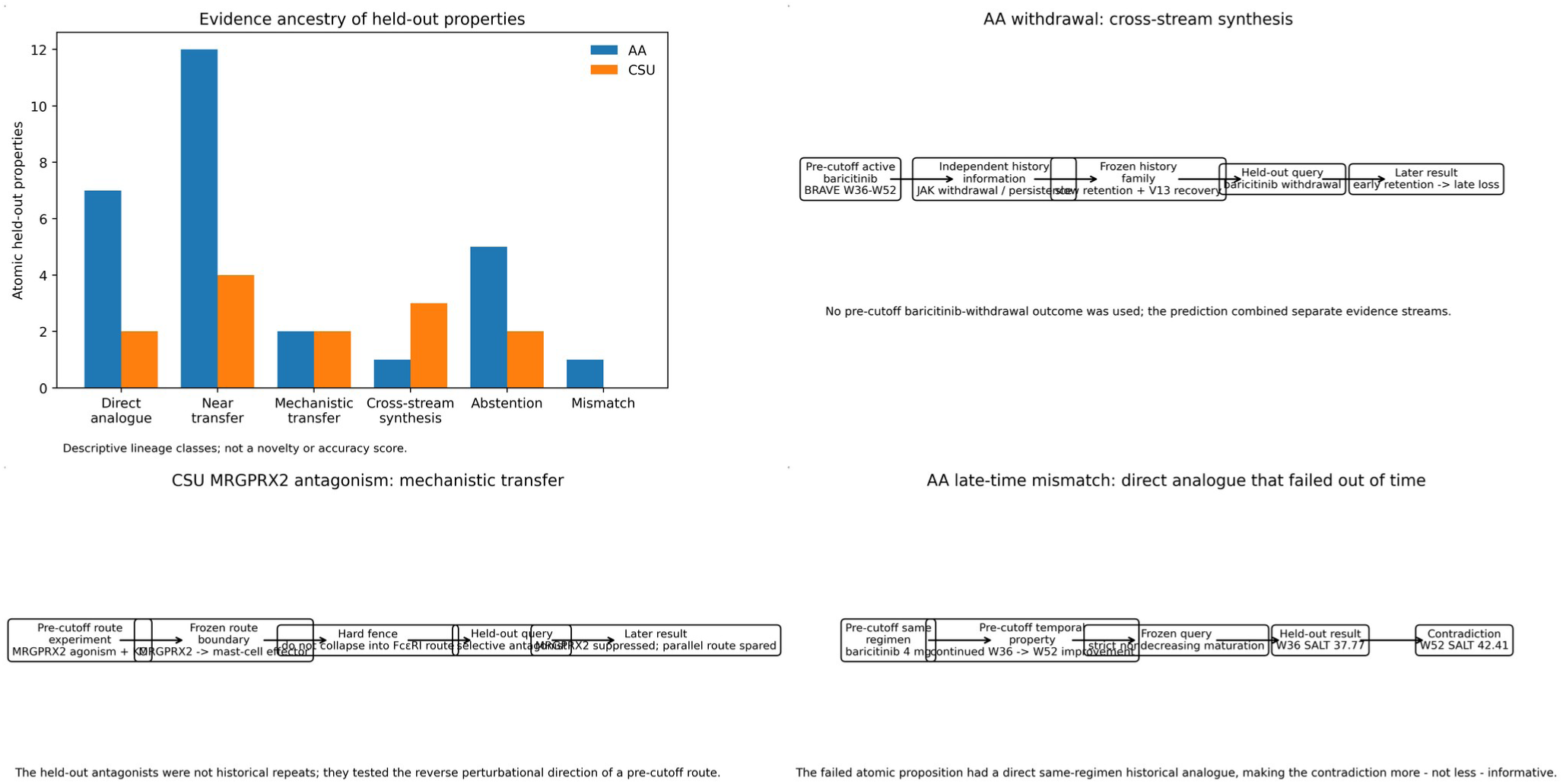
Evidence ancestry of held-out predictions. a, Atomic held-out properties separated by evidence-lineage class. b, Baricitinib withdrawal required cross-stream synthesis because no pre-cutoff baricitinib-withdrawal result was used. c, MRGPRX2 antagonism is mechanistic transfer from route-separation evidence. d, The AA week-36-to-52 mismatch contradicted a direct same-regimen historical temporal property. Counts are descriptive and are not a novelty or accuracy score.

### AA tests persistence, temporal order, route and information boundaries

AA provided the broader primary validation set: 21 primary studies across 15 dependency families, plus two sensitivity studies. Study-level scoring yielded 14 concordant, four concordant with a prespecified abstention boundary, three partially concordant and none labelled fully discordant; dependency adjustment yielded 12 concordant and three partially concordant families. These are audit counts over heterogeneous information properties, not a single predictive-accuracy estimate (Supplementary Note 8; Supplementary Table S7).

Withdrawal was the clearest high-information test. The frozen baricitinib-withdrawal trajectories retained substantial latent benefit early and then decayed progressively. In BRAVE-AA1, no withdrawn responder had lost treatment benefit at four weeks, about 10-11% had lost benefit by eight weeks, and approximately 80% had lost benefit by 100 weeks after withdrawal, while continued treatment largely maintained response.^18^ The model and clinical axes are deliberately shown separately because latent retention is not a patient relapse probability (AA-PUB-003; Supplementary Note 8; Supplementary Tables S7-S9).

Several ritlecitinib and baricitinib studies supported nondecreasing maturation over weeks to months, but the lineage audit showed that these tests were not equally novel: some repeated direct historical treatment-time properties, whereas others extended them to new times or cohorts. The late baricitinib real-world trajectory above was the genuine exception. At study level it remains partially concordant because the earlier active-treatment trajectory was compatible; at atomic level, the registered week-36-to-52 monotonicity component is discordant (AA-PUB-099; Supplementary Note 8; Supplementary Table S8, row AA-PUB-099.P1).

Route and history tests further separated what was identifiable from what was not. Post-cutoff topical tofacitinib supported the frozen local-JAK route without establishing systemic equivalence. By contrast, nominal baricitinib and ritlecitinib doses could not be converted into a mechanistic potency rank, and observational prior-JAK subgroups did not identify causal history effects. These withheld comparisons count as validation of information boundaries, not predictions of equality (Supplementary Note 8; Supplementary Tables S7-S9).

**Figure 4.**
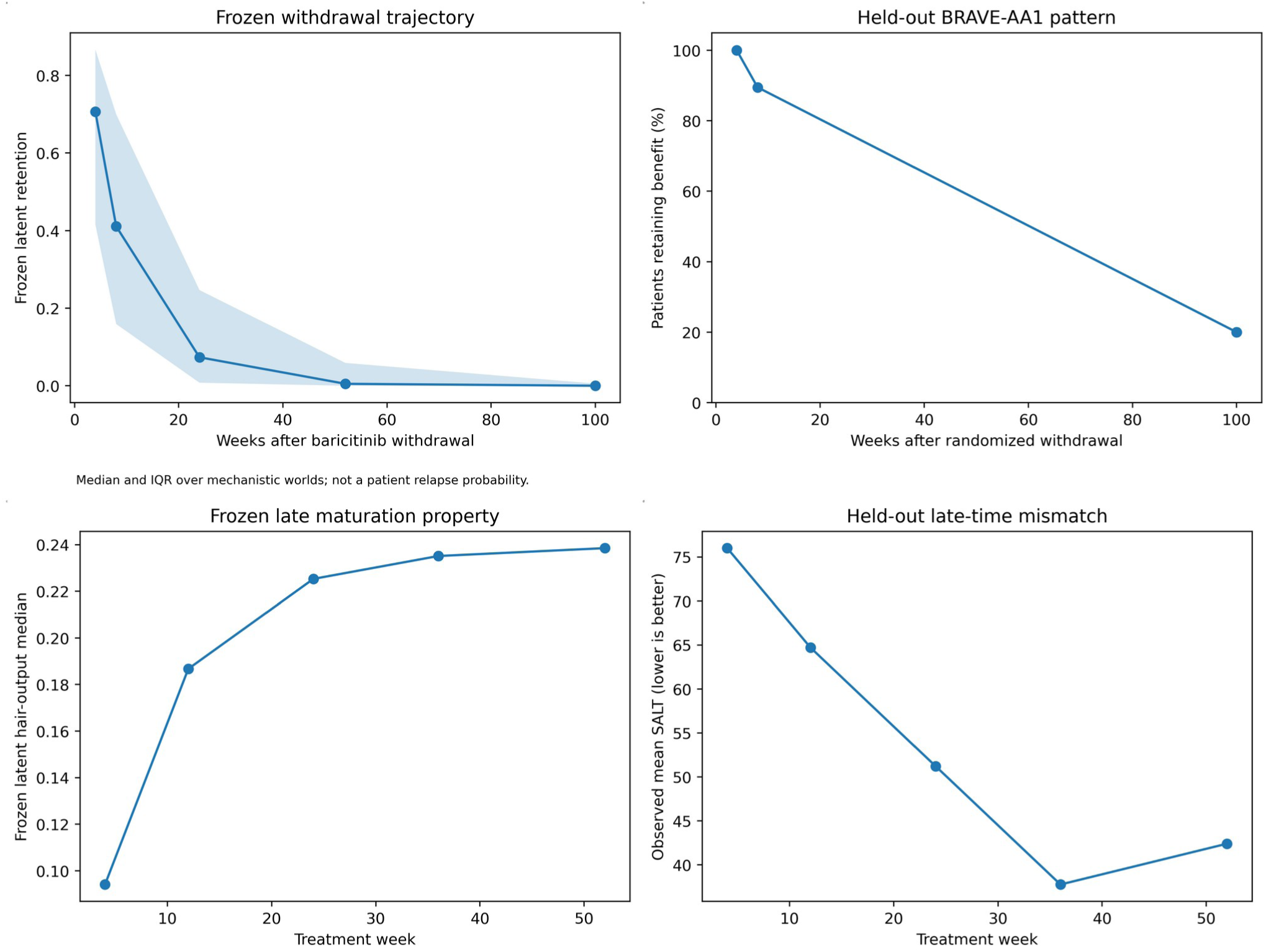
High-information validation in AA. a,b, Frozen withdrawal persistence and held-out BRAVE-AA1 are shown on separate latent and patient-level axes. c,d, The frozen late-maturation property and AA-PUB-099 outcome expose the week-36-to-52 contradiction. Full AA results and lineage are in Supplementary Notes 8 and 10 and Supplementary Tables S7-S9.

### CSU has narrow primary clinical coverage but strong route and timing tests

The CSU primary clinical register was smaller and narrower: five primary studies, all using omalizumab 300 mg. Three were concordant and two were partially concordant because the public reports did not expose all intermediate values required for the prespecified temporal order. This anti-IgE primary layer therefore supports the historical clinical bridge but should not be read as broad treatment-class validation. Five sensitivity tests supplied greater mechanistic diversity (Supplementary Note 9; Supplementary Table S7).

Cyclosporine supplied a rapid timing test. The frozen system predicted improvement during the fixed one-to-two-week active phase. In the held-out cohort, mean UAS7 fell from 31 at baseline to 5.78 after one week and 3.56 after two weeks.^20^ The lineage was near transfer rather than direct replay: pre-cutoff cyclosporine studies established clinical and calcineurin-route effects, but not this exact day-7-to-day-14 UAS7 ordering (CSU-PUB-089; Supplementary Note 9; Supplementary Tables S8-S9).

**Figure 5.**
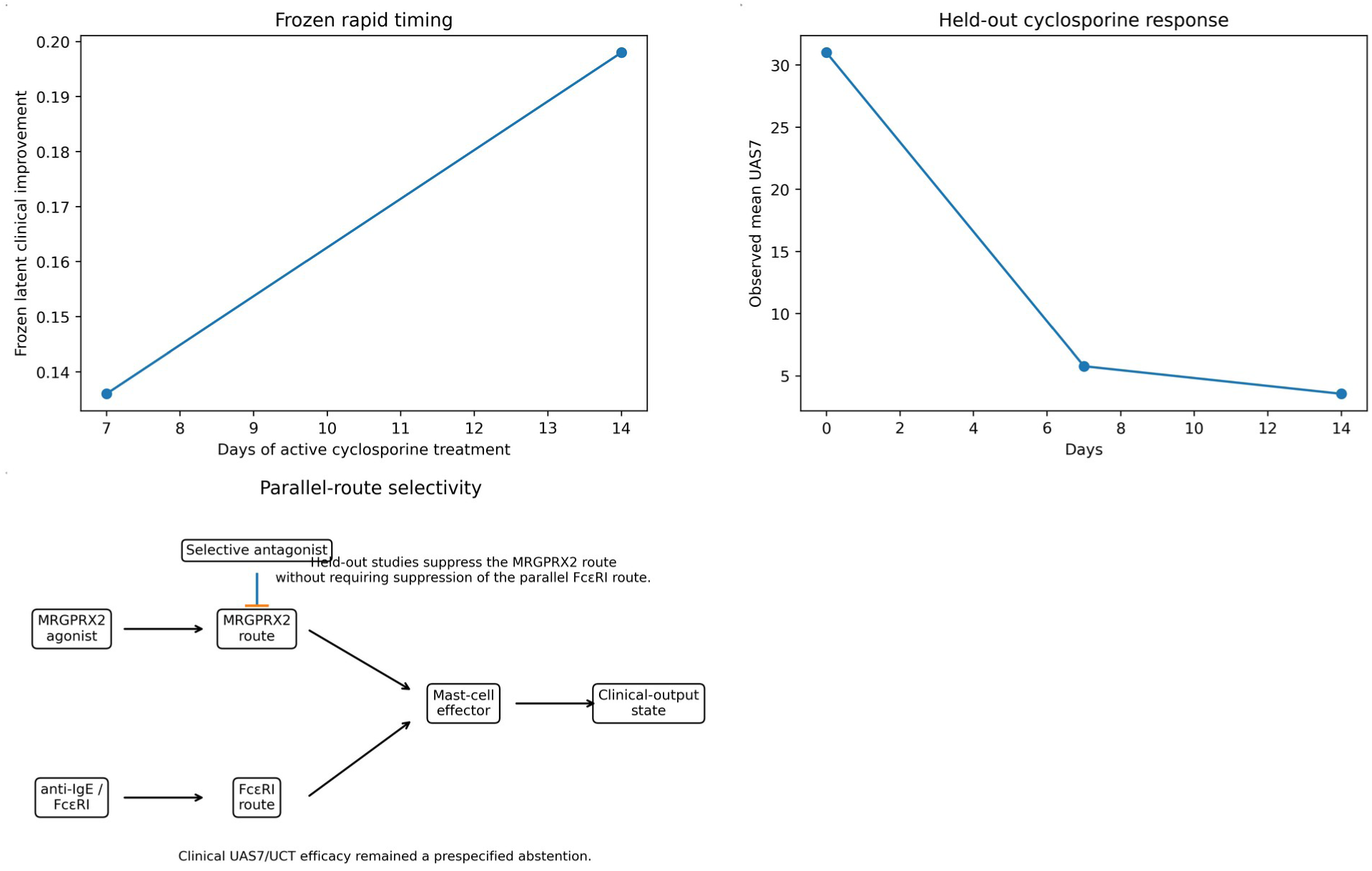
High-information validation in CSU. a,b, Rapid cyclosporine timing is shown on separate latent and UAS7 axes. c, Selective MRGPRX2 perturbation tests a route represented separately from Fc-epsilon-RI signalling; patient-level clinical efficacy remains outside the identified observation resolution. Complete CSU scoring is in Supplementary Note 9.

The two MRGPRX2 studies were higher-information route tests. Selective antagonists inhibited MRGPRX2-driven mast-cell degranulation in human cellular and skin systems, and one experiment directly preserved the anti-IgE/Fc-epsilon-RI component while blocking the Substance-P/MRGPRX2 component.^21,22^ The frozen system had encoded those routes separately before outcome reveal. It did not, however, claim a patient UAS7 or UCT effect from the mechanistic assays; that clinical extrapolation remained an explicit abstention (CSU-PUB-127/128; Supplementary Note 9; Supplementary Tables S8-S9).

### Finite ensembles preserve the registered uncertainty with auditable numerical error

The robustness question is whether numerical representation changes the registered scientific conclusions, not whether the finite ensemble resembles a patient population. Under the prespecified evidence-to-q adapter strengths alpha=1.2, 1.6 and 2.0, the favoured side of the registered q controllers did not reverse; allocation strength changed, but the qualitative evidence direction remained stable (Supplementary Note 11; Supplementary Table S12).

The AA candidate allocation had weighted effective sample size (ESS) 40.1%, maximum final q error 0.00546, 16,017 unique resampled candidates and 13,599 distinct structural signatures at N=16,384. CSU had weighted ESS 99.6%, maximum final q error 0.00331 and 16,533 distinct structural signatures at N=32,768. For CSU, N=16,384 failed the prespecified three-seed standardized-output stability rule (maximum difference 0.01893); N=32,768 was the smallest passing tier (0.00394). Function/time/history censuses confirmed broad representation of linear, Hill-like, threshold, power, gating, delay/persistence and multiple history families, but these frequencies are coverage diagnostics, not biological prevalence (Supplementary Notes 5 and 11; Supplementary Tables S13-S15).

**Figure 6.**
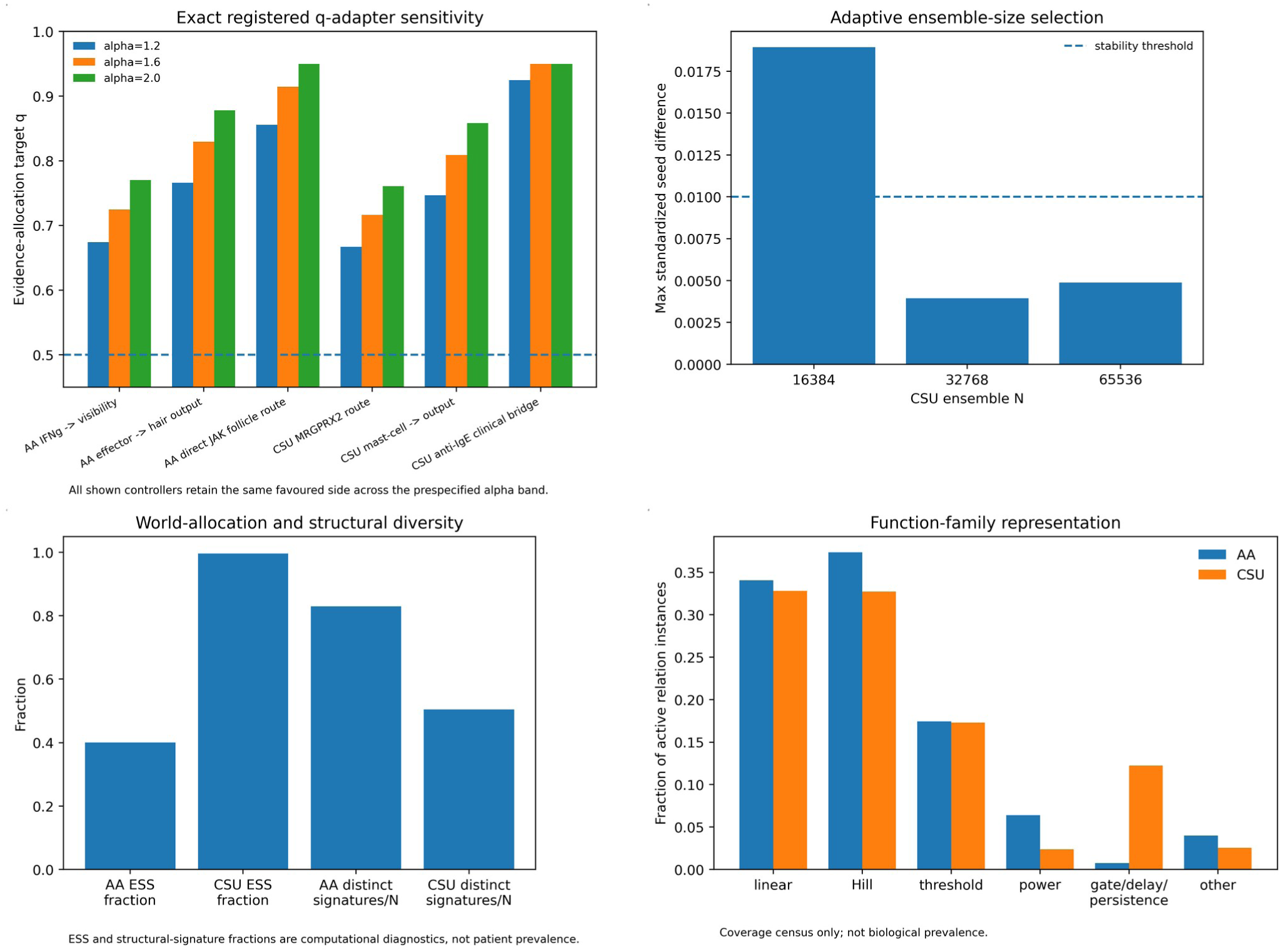
Robustness and coverage audits. a, Exact registered q-controller values across the prespecified evidence-adapter strengths alpha=1.2, 1.6 and 2.0; the favoured side does not reverse in the shown controllers. b, CSU ensemble size selected by a multiseed standardized-output gate. c, Allocation ESS and distinct structural-signature coverage. d, Function-family representation. These are computational diagnostics, not patient or biological prevalence.

### A separate decision layer supports research perturbation prioritization

The frozen ensemble can be used downstream to compare researcher-declared biological objectives. The current research interface searches package-declared molecule/regimen inputs and optional mechanistic perturbations on matched worlds; objective-fit percentages summarize decision-layer loss rather than potency, efficacy or response probability. The full demonstration is moved to Extended Data Fig. 2 because it is an application of the validated ensemble rather than the primary scientific result. An interactive research perturbation interface will be provided at https://gen-lang-client-0942684525.uc.r.appspot.com/. It is not clinical decision support, dose optimization, safety evaluation or patient-specific treatment advice.

## Discussion

This study tests a methodological premise: heterogeneous biomedical knowledge can contain executable mechanistic information before it contains the commensurate measurements required to fit one system model. ECMS makes that information explicit by separating information type from evidence strength, restricting ontology and relation semantics to what is identified, and sampling whole mechanistic worlds rather than silently filling unknown dimensions with one expert model.

The lineage audit is central to interpreting the historical results and provides a deliberately conservative internal baseline. Many simple positive-treatment findings were direct or near historical analogues; they are useful consistency checks but weak evidence for a complex framework and could plausibly be recovered by nearest-historical matching. The more informative examples were those in which later experiments interrogated a different aspect of the pre-cutoff information state: withdrawal assembled from active-treatment and history streams, MRGPRX2 antagonism transferred from route-dissection experiments, dual UAS7/UCT observation semantics combined independent clinical and measurement streams, and abstention preserved causal ignorance. Conversely, the AA late-time mismatch remained visible even though its contradicted temporal property had strong historical ancestry. The present study does not include a formally matched suite of alternative computational models; such benchmarking should be performed prospectively or on a predeclared common benchmark rather than reconstructed after seeing this validation set.

ECMS occupies a different point in the modelling landscape from several active approaches. Biomedical knowledge graphs such as PrimeKG organize large heterogeneous relation sets, and graph-neural drug models such as TxGNN learn predictive representations from them.^11,16^ Perturbation models such as GEARS, CellOT and scGPT learn intervention-response mappings from high-dimensional experimental data, while CINEMA-OT brings explicit causal matching to single-cell perturbation analysis.^12,13,14,15^ Deep structural causal models instead assume or learn a causal generative structure suitable for interventional and counterfactual inference.^17,23^ Automated assembly systems such as INDRA convert textual mechanistic statements into normalized executable representations.^9,10^ ECMS is complementary to each: its scientific object is an evidence-constrained possibility space for non-commensurate pre-calibration literature, with explicit resolution fences and first-class abstention. A neutral architecture comparison, with claim-specific primary references, is in Supplementary Note 12 and Supplementary Table S6.

ECMS is also complementary rather than competitive with calibrated mechanistic systems modelling and QSP. Once matched exposure, target occupancy, quantitative biomarkers, observation maps and clinical outcomes are sufficient to identify the quantities relevant to a decision, a calibrated model is preferable for those quantities.^3,4^ The ECMS evidence atlas and possibility-space semantics can then serve as a structured precursor: qualitative relations can be promoted to quantitative modules as calibration becomes defensible.

Several limitations bound the present evidence. The historical reconstruction was performed retrospectively in 2026 and therefore cannot create investigator naivety equivalent to prospective preregistration. The LLM-assisted semantic-compilation log, pre-cutoff source restrictions, human source adjudication and checksum firewalls make the process auditable and reduce leakage risk, but they cannot prove that retrospective bias was impossible. The disease atlases are auditable representative corpora rather than exhaustive systematic reviews. CSU primary clinical coverage is narrow and omalizumab-dominated. Several temporal tests are reporting-limited. The evidence-to-q adapter is an engineering mapping that combines design, transportability, statistical and event-match weights; the current alpha=1.2/1.6/2.0 sensitivity band established stability of controller direction but is not a comprehensive prior-sensitivity analysis. Broader sensitivity to alternative reference measures, caps and leave-one-dependency-out constructions remains future work. The finite function and history grammar is broad but declared, not unlimited; an observation outside it is a legitimate falsification rather than permission for retrospective expansion. Finally, epistemic world frequencies are not patient heterogeneity, and latent outputs are not calibrated clinical scores.

The decisive next test is prospective: construct H(E), freeze a new disease or target programme before the discriminating experiment is known, use the lineage audit to state exactly why each perturbational proposition is expected, and then expose the result. If successful, ECMS could support evidence-constrained virtual screening of biological interventions before molecule screening; if it fails, the failed information object should identify which scientific coordinate - ontology, route, function, time/history, input or observation map - requires revision in H(E_new).

## Methods

### Study design and governance

We performed a two-disease retrospective outcome-blind historical reconstruction. The construction evidence state was restricted to information publicly available on or before 31 December 2023. Earliest public availability, not publication year alone, governed eligibility. Concepts, ontology bridges, biological intuition and external reasoning capable of changing the possibility space obeyed the same cutoff. General ECMS mathematics is not repeated here; the authoritative reader-facing definition is Supplementary Methodology (Supplementary Note 1).

### LLM-assisted semantic compilation and governance

From 1 to 20 August 2026, package construction used OpenAI GPT 6-sol thinking models at high reasoning effort as iterative semantic-compilation and biological-intuition aids. For construction decisions, the models were supplied with pre-cutoff source material and the prespecified ECMS information-anatomy rules and could propose source-grounded ontology bridges, information-class translations, relation/boundary alternatives and function/time/history candidates. They were not authorized to treat their own output as evidence, generate unsupported clinical-response magnitudes, infer cross-molecule engagement from nominal dose, or make final scientific acceptance decisions. Investigators manually verified proposals against the source material and adjudicated accept/revise/reject decisions; the final frozen package and a manuscript-facing biological-intuition log were manually checked. The process was iterative rather than zero-shot. Individual prompt transcripts are not part of the public scientific package because prompts served as workflow scaffolding rather than evidence; partial prompt traces can be provided to reviewers if specifically informative. The log records accepted frozen rules, linked pre-cutoff evidence identifiers, model family, execution window, prohibitions and human adjudication (Supplementary Table S18).

### Evidence objects and information anatomy

Primary mechanistic, interventional, observational, omics and pharmacological sources were decomposed into atomic findings with provenance, context, perturbation/exposure, readout, uncertainty where available, dependency grouping, transportability and contradiction/context flags. Each finding was classified by the strongest directly supported information type: node/set relevance, topology/possibility boundary, direction/order, bound, function family, time/history or observation map. Multiple dependent findings were aggregated within source/dependency programmes before independent evidence programmes contributed to soft allocation.

### Possibility-space construction and evidence allocation

Named biological coordinates were treated as unresolved sets unless evidence showed that a split altered executable consequences. Relationship envelopes were registered as bidirectional possibility boundaries whose directed realization, latent completion, function, context and time/history family varied by world. Hard fences removed incompatible worlds. For a soft event C_j, dependency-aware evidence compilation produced target allocation q_j relative to a declared reference measure. Candidate-world weights were adjusted by minimum-information projection to approach registered margins while preserving hard fences. q is an ensemble-construction controller, not a posterior probability.

### Hierarchical world generation and adaptive ensemble size

Worlds were generated structural first: context and set membership; boundary and latent-subgraph realization; overlap; function/sign; time/history family; numerical execution values; perturbation inputs; observation maps. AA and CSU each began from 100,000 candidate worlds. Frozen N was the smallest audited tier meeting q-error, hard-fence, multiseed and standardized benchmark-output stability criteria. Detailed q tables, ESS, structural signatures, function/time censuses and N-selection diagnostics are in Supplementary Notes 4-5 and 11.

### Input and observation maps

Interventions were mapped through molecule/regimen-specific frozen input maps. Nominal doses across different molecules were not interpreted as a common target-engagement scale. Observation maps linked latent variables to the qualitative or ordinal resolution justified by historical evidence. No numerical transformation from latent AA hair-output to SALT, or from latent CSU activity to UAS7/UCT, was assumed.

### Validation freeze and concordance grading

Post-cutoff intervention identity, route, regimen, evaluable times, observation semantics and information class were frozen before downstream outcomes were entered. Outcome-blind prediction artifacts were checksummed before held-out outcome extraction. Each registered atomic property was scored as concordant, concordant with an abstention boundary, partially concordant or discordant. Partial results were further classified as true model-data mismatch, reporting insufficiency, input non-identifiability or observation non-identifiability. Study-level grades summarize registered atomic components but do not override an explicitly discordant atomic proposition. The complete canonical table is Supplementary Table S7 and the atomic table is Supplementary Table S8.

### Evidence-to-prediction lineage audit

For every held-out atomic proposition we backtraced the frozen validation query through input and observation maps to the world-level q events and hard fences used by that proposition, then to pre-cutoff atomic evidence identifiers, dependency programmes and original source identities. We recorded whether the same molecule, regimen, endpoint, time/property and exact study/trial programme were already represented. Lineages were classified descriptively as direct historical analogue, near transfer, mechanistic transfer, cross-stream synthesis, abstention boundary or mismatch. No numerical novelty score was used and post-cutoff knowledge was not used to infer the lineage. Full records are Supplementary Note 10 and Supplementary Table S9.

### Statistical and numerical analyses

Validation counts are descriptive. No pooled treatment-response accuracy was computed across heterogeneous information classes, and prespecified abstentions were not counted as empirical successes. At atomic-property level we report 29 empirically concordant answered propositions, one mismatch, four committed propositions that could not be fully assessed, and seven prespecified abstentions. Dependency-family summaries reduce pseudo-replication. Numerical audits report candidate-world weighted effective sample size, final q error, hard-fence failures, distinct structural signatures and multiseed standardized benchmark-output differences. q-adapter sensitivity used the prespecified alpha values 1.2, 1.6 and 2.0 on the same scientific grammar; it is an allocation-target sensitivity analysis rather than a confidence interval.

### Code and data availability

The reproducible code/data archive and scoring scripts will be released through the study GitHub repository at https://[ECMS-GITHUB-URL-TO-BE-ADDED] before submission/public release. The interactive research perturbation/decision interface will be available at https://gen-lang-client-0942684525.uc.r.appspot.com/.

The interface executes the manuscript’s frozen/current ECMS perturbation semantics for research exploration only and is not clinical decision support. Supplementary Software 1 contains the manuscript-matched package archive. File identities and checksums are listed in Supplementary Note 13.

## Extended Data

**Extended Data Figure 1.**
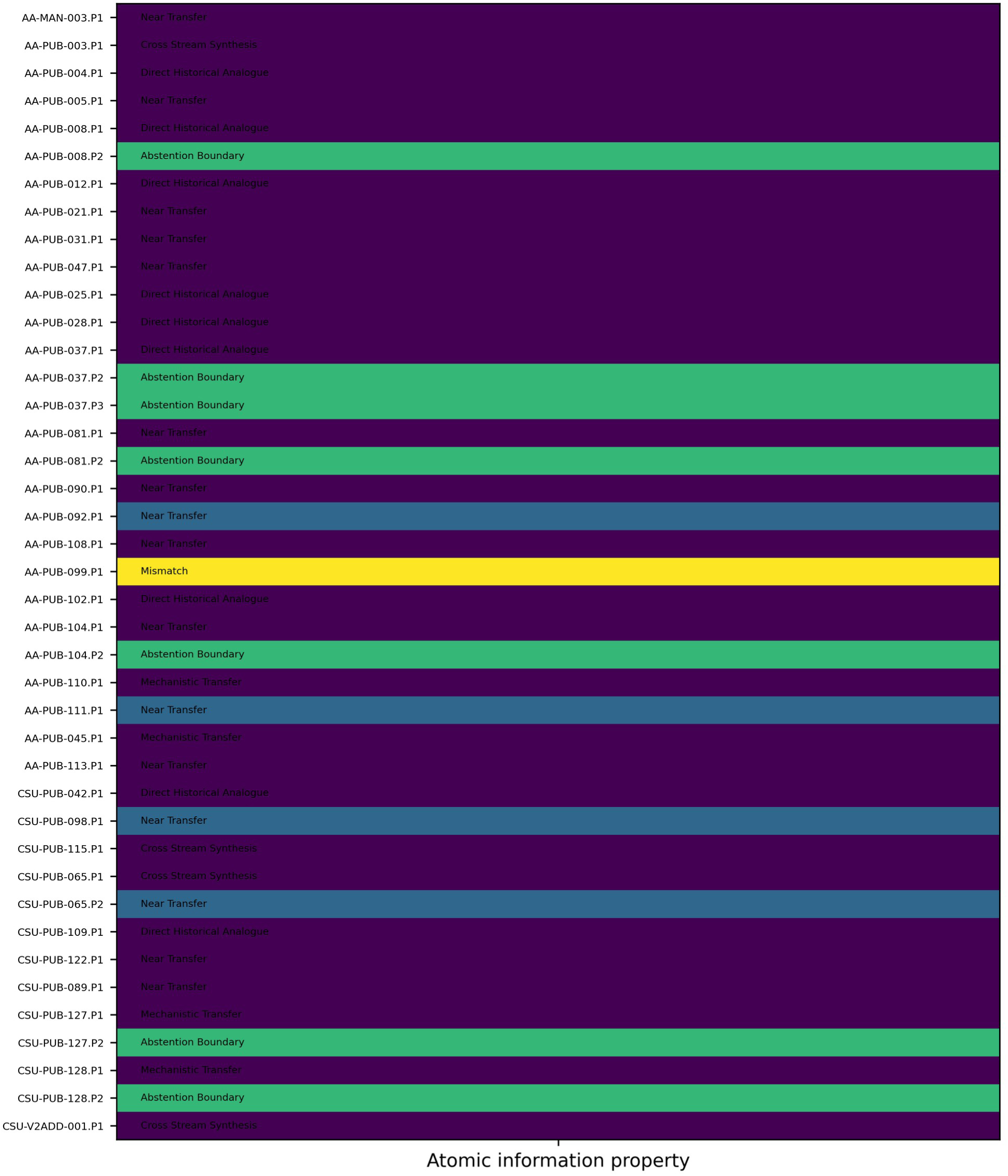
Complete atomic information-property landscape. Each registered property is shown once with its grade and evidence-lineage class. The underlying canonical records are Supplementary Tables S8-S9.

**Extended Data Figure 2.**
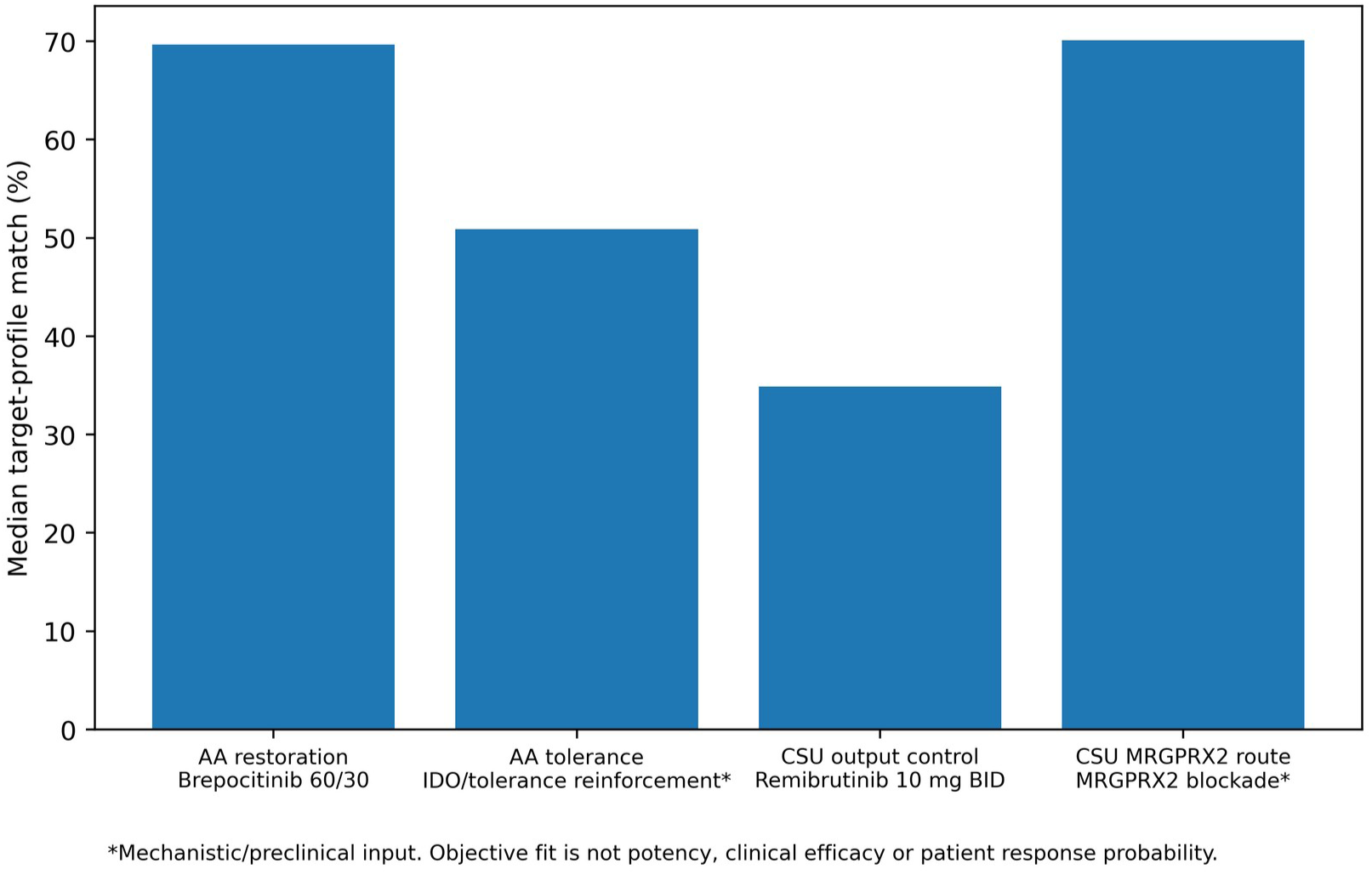
Objective-dependent perturbation prioritization on unchanged ensembles. The online interface is a research exploration tool; target-profile match is not potency, clinical efficacy or a patient-response probability.

## Supplementary Methodology (Supplementary Note 1)

Evidence-Constrained Mechanistic Synthesis: general methodology and mathematical specification | 25 August 2026

|  |  |
| --- | --- |
| Purpose | A rigorous, disease-agnostic description of how ECMS converts heterogeneous biomedical evidence into an auditable ensemble of mechanistic worlds. |
| Scientific regime | Pre-calibration: evidence is rich enough to constrain mechanisms but insufficient to justify one fully calibrated quantitative system model. |
| Fundamental object | An evidence-constrained possibility space of complete mechanistic worlds, not a single graph and not a fixed list of expert models. |
| Reference policy | Only public methodological/statistical references are cited here. Disease-specific historical sources belong to disease implementations and validation artifacts, not to the definition of ECMS. |

**Central rule: represent no more biological detail, causal direction, functional form, numerical precision or temporal structure than the evidence actually identifies. Everything else remains an explicit possibility.**

## 1. Why ECMS is needed: the pre-calibration problem

Mechanistic research frequently reaches a stage at which the literature contains enough information to exclude many biological explanations and to support useful perturbation reasoning, but not enough homogeneous quantitative data to identify one calibrated model. At this stage, forcing every finding into a coefficient creates precision that the source never contained, whereas waiting for complete calibration discards real mechanistic information. ECMS is designed for this intermediate regime. It is complementary to quantitative systems pharmacology (QSP) and other calibrated mechanistic models: when calibration-grade data exist for the question and context of use, a calibrated model is generally preferable [10,11].

The method starts from a distinction that is easy to state but consequential in practice: different findings answer different mathematical questions. An intervention may establish that one process can influence another without identifying the mediator. A comparative experiment may establish A>B without identifying the numerical difference. A time-course may identify onset before week 4 and persistence after withdrawal without identifying an exponential half-life. An association may identify shared information without identifying causal direction. ECMS preserves those differences instead of collapsing all findings into one scalar evidence score or one parameter-fitting objective.

The resulting object is not a single “best” pathway. ECMS defines a possibility space of complete mechanistic worlds. Each world is a fully executable assignment of biological context, active members of unresolved sets, directed structural realization inside possibility boundaries, function classes, numerical execution values, temporal/history behavior, perturbation-input mapping and observation mapping. Evidence constrains which worlds are admissible and how frequently evidence-supported events are represented. The finite ensemble used for computation is an auditable sample of that constrained space, not the biological universe itself.

### 1.1 Three uncertainties that ECMS keeps separate

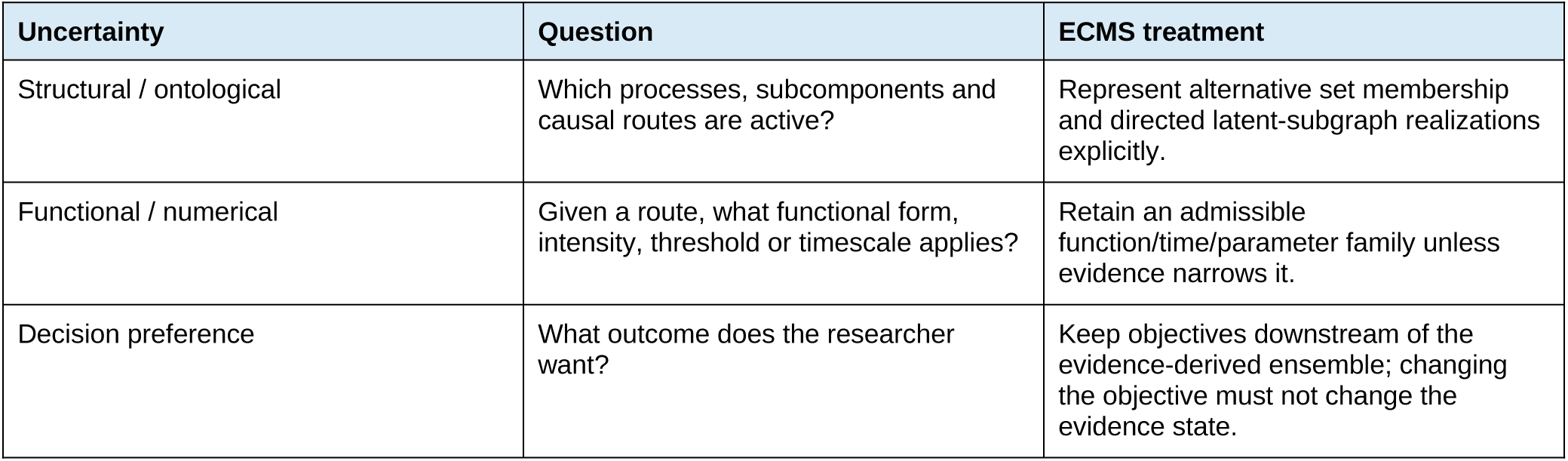

| Uncertainty | Question | ECMS treatment |
| --- | --- | --- |
| Structural / ontological | Which processes, subcomponents and causal routes are active? | Represent alternative set membership and directed latent-subgraph realizations explicitly. |
| Functional / numerical | Given a route, what functional form, intensity, threshold or timescale applies? | Retain an admissible function/time/parameter family unless evidence narrows it. |
| Decision preference | What outcome does the researcher want? | Keep objectives downstream of the evidence-derived ensemble; changing the objective must not change the evidence state. |

Patient-to-patient heterogeneity is a fourth concept and must not be silently identified with any of the three. An ECMS ensemble is epistemic unless a separate population model has been constructed. Therefore an ensemble frequency is not automatically a patient prevalence or response probability.

## 2. Core philosophy: information before parameters

### 2.1 Minimum commitment

The minimum-commitment rule is the organizing principle of ECMS: encode the weakest mathematical statement that reproduces the information actually justified by a finding. Deeper information is never inferred merely because a shallower statement is known. For example, evidence for a possible influence permits a structural relation; it does not by itself identify sign, magnitude, function shape, timing or observation calibration. This is analogous to maximum-entropy and minimum-discrimination-information reasoning, in which the representation is changed only as much as required by stated constraints [1–4].

### 2.2 Information content as uncertainty reduction

Information content and evidence reliability are distinct. Information content asks how restrictive a mathematical statement would be if accepted. Reliability asks how strongly the evidence supports assigning that restriction to the mechanistic ensemble. ECMS uses bits as a transparent audit unit for the first question.

Let the reference state space be Ω_0_ and let a hard constraint C retain the subset Ω_C. Under a uniform reference measure, the information supplied by the hard restriction is

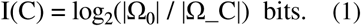

For a non-uniform reference distribution P_0_, the same quantity is

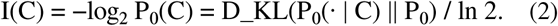

Consider the deliberately simple example used to make the principle tangible. Let A and B each take integer values 1,…,10, so that Ω_0_ contains 100 equally plausible ordered pairs. The statement A>B retains 45 pairs and contributes log₂(100/45)=1.152 bits. Requiring A−B≥3 retains 28 pairs and contributes 1.837 bits. Fixing A=5 and B=7 retains one pair and contributes 6.644 bits. The point is not that exact values are always better; the point is that they are more restrictive and therefore require correspondingly stronger evidence.

**Figure 1.**
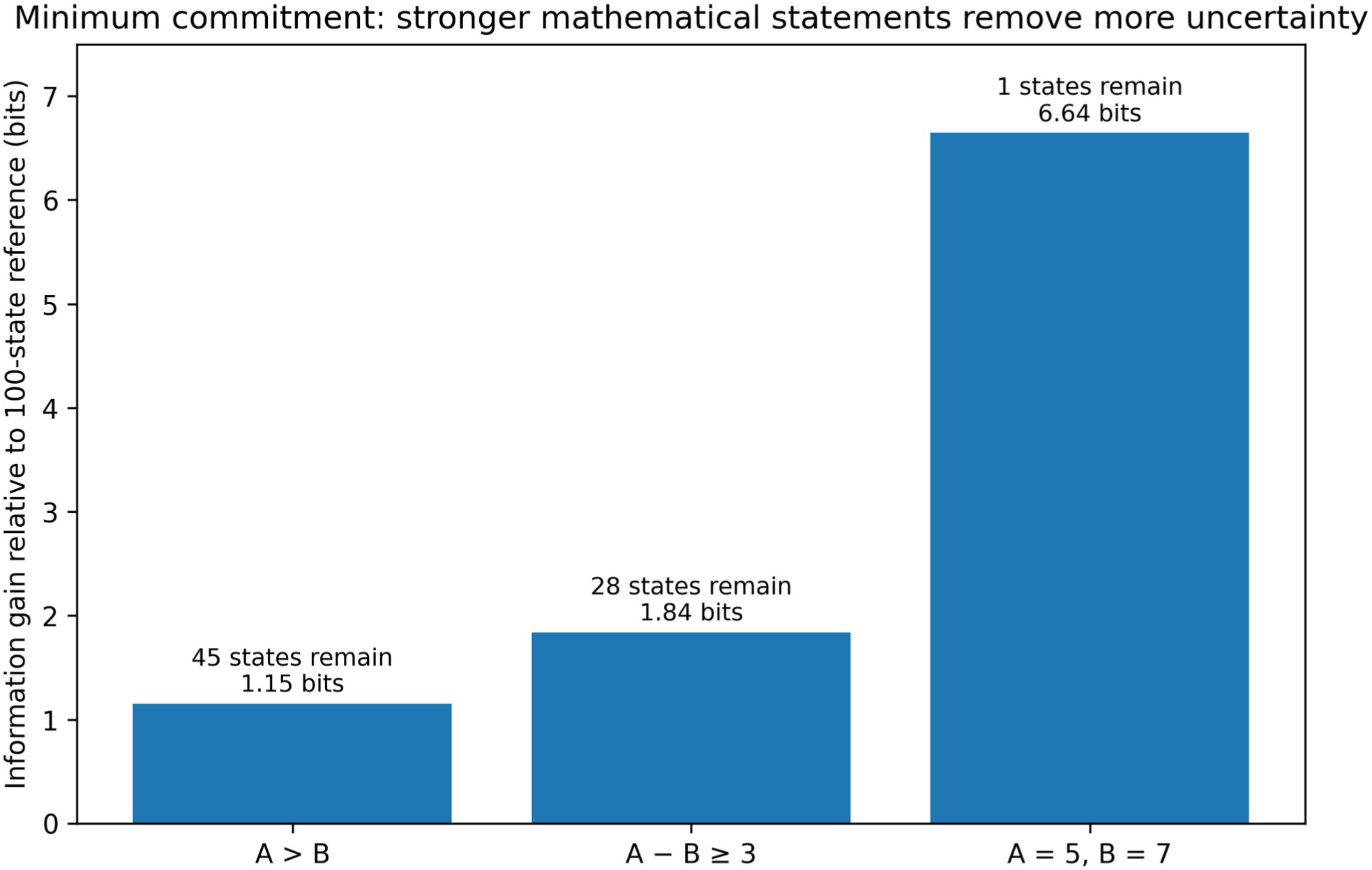
Information content as uncertainty reduction. The toy 10×10 state space shows why an ordinal statement should not be silently upgraded to a numerical gap or exact value. The counting formulation is a special case of Shannon/Kullback-Leibler information [1,2].

### 2.3 Soft evidence and information relative to a reference event frequency

Many biological claims are not hard facts. If event C has reference allocation p_0_ but compiled evidence targets allocation q, all worlds need not be deleted; the distribution can instead be reweighted. For a single binary event, the information added relative to the reference is the Bernoulli Kullback-Leibler divergence

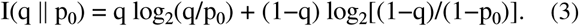

For example, moving a symmetric reference event from p_0_=0.5 to q=0.70 adds only 0.119 bits. This illustrates why q should not be read as a biological truth probability. q is a construction target describing how the evidence changes allocation relative to a declared reference measure.

With multiple overlapping constraints, marginal information values are generally not additive. The correct global audit is the relative entropy between the final least-committal world distribution Q* and the reference measure μ_0_:

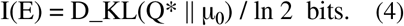

This prevents double-counting information when several evidence objects constrain the same worlds.

### 2.4 Information anatomy

| Information type | What the evidence can say | Mathematical object | What it does not license |
| --- | --- | --- | --- |
| Representation | The object/process must be represented. | Membership in an unresolved set or context variable. | No causal relation. |
| Topology / influence | A route can connect two biological regions. | A possibility boundary and admissible path event. | Mediator identity, sign or magnitude. |
| Direction / sign | Increasing a source moves an outcome up or down in a context. | Sign or directional event on a realized path. | Numerical gain or universality. |
| Order / comparison | $A > B$, $A > B > C$, or event A precedes B. | Joint world-level comparison event. | A numerical gap. |
| Bound / ceiling / gap | A quantity lies in an interval or exceeds a minimum separation. | Inequality or interval constraint. | Arbitrary function fitting. |
| Function class | Monotone, saturating, threshold, reversing, hysteretic, persistent, etc. | Restriction of the admissible function family. | Exact parameters unless measured. |
| Magnitude | A transportable effect/range is measured. | Numerical parameter/bound distribution. | Transfer to incompatible contexts. |
| Time / history | Onset, lag, order, persistence, recovery or memory is supported. | Clock/history-family restriction. | A smooth curve or half-life not measured. |
| Observation map | A latent state can be related to a measured endpoint. | Mapping at the supported resolution. | Causal status of the biomarker. |
| Context / counterexample | The relation differs by tissue, stage, endotype or other context. | Context gate or structural alternative. | Averaging away the difference. |
| Combination / interaction | Joint action is measured relative to a reference. | Interaction constraint. | Synergy merely because two perturbations are combined. |

The information ladder is not a claim that every study should progress through all levels. Stopping at topology or order is often the scientifically correct result. Unused dimensions remain uncertainty to be sampled, not blanks to be filled by convenience.

## 3. Formal objects and notation

### 3.1 Evidence state and constrained world space

Let E denote the evidence state used for a particular ECMS release. E includes atomic findings, provenance, dependency structure, context and transportability labels, hard fences and soft allocation events. Let ℋ_o_ be the declared pre-evidence grammar of executable worlds for the scientific question. Hard evidence fences F_k define the admissible support

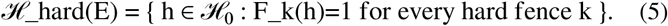

A reference measure μ _o_is then declared over ℋ_hard(E). μ _o_is an engineering measure over residual uncertainty, not a prior probability of biological truth. Soft evidence defines target margins on events C_j(h). The shorthand H(E) refers to the resulting evidence-constrained possibility space: its admissible support, its registered soft constraints and the reference measure required to sample it reproducibly.

### 3.2 Biological variables are unresolved sets

A named ECMS variable is not assumed to be an indivisible biological entity. It is an unresolved set of admissible members or substates:

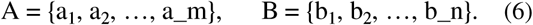

The purpose of the set is compression without false certainty. In a world h, active membership can be represented by M_A^(h)⊆A and M_B^(h)⊆B. A split into explicit variables is required only when the evidence demonstrates behaviorally different consequences that can change an executable prediction. Naming two biological subtypes is not sufficient reason to split them.

A useful invariance test is permutation equivalence: if two proposed members can be exchanged in every admissible world without changing any relation event, perturbation response, comparison, temporal behavior or observation, the distinction contributes no ECMS information and should remain compressed.

### 3.3 The bidirectional A ↔ X ↔ B possibility boundary

The reader-facing ECMS representation of an unresolved biological relationship is a bidirectional possibility boundary

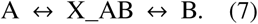

This notation is intentionally different from a conventional bidirectional causal edge. X_AB is not one hidden node. It denotes a family X_AB of admissible latent subgraph completions connecting the two unresolved sets. The double arrows indicate that the boundary exposes interfaces in both directions to the world generator. They do not state that both causal directions are active in every world.

For a boundary between A and B, define the candidate interface set

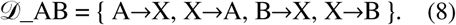

A complete world h selects an active subset D_AB^(h)⊆D_AB, an internal latent realization X_AB^(h)∈X_AB, and any evidence-admissible active members of A and B. Evidence and hard fences determine which subsets are allowed and how frequently particular realized path events occur. Consequently, the same boundary can generate a forward influence world, a reverse influence world, a common-cause/association world, a feedback world, or a more complex partially resolved realization. A direct resolved relation is the limiting case in which the latent completion collapses to an identity/zero-depth mediator at the modelled resolution.

This boundary formalism is the central device by which ECMS avoids making ontology or direction more precise than the evidence. If causal evidence supports A→B, the corresponding event is defined on a complete world as the existence of an admissible directed path from an active member of A to an active member of B. Reverse influence is not automatically required merely because the possibility boundary is bidirectional. Conversely, if feedback is supported, both directed path events can be active in the same world. Pure association can be represented by X→A and X→B without a directed path A→B or B→A [8].

**Figure 2.**
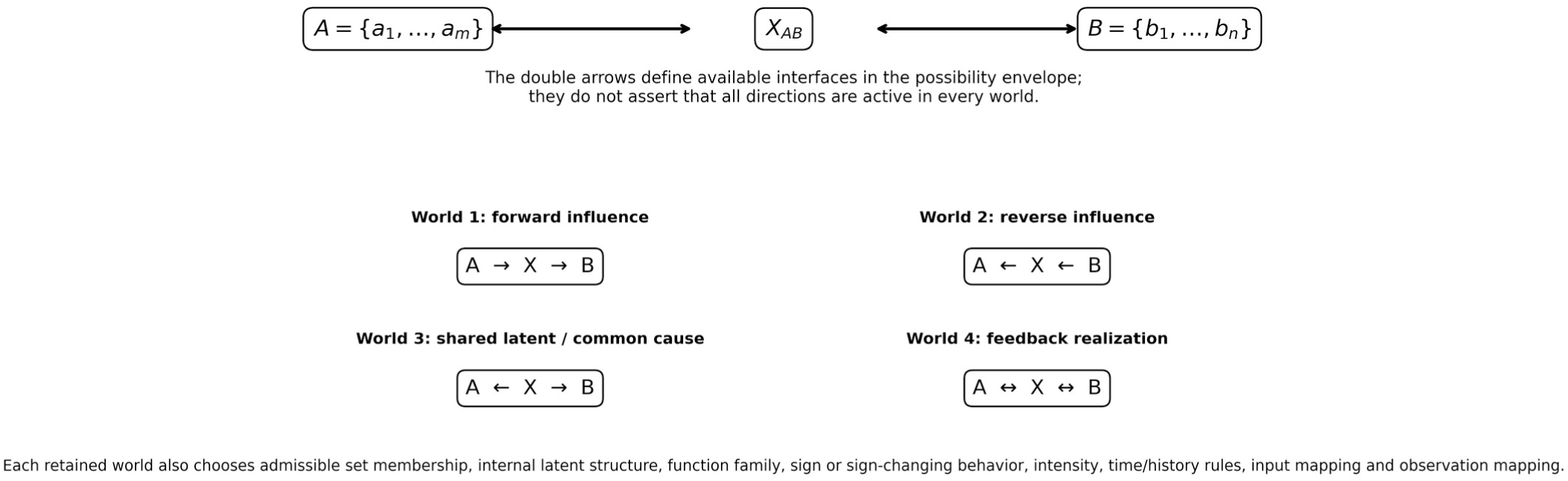
Bidirectional possibility boundary. A↔X_AB↔B is a possibility envelope. Each mechanistic world instantiates an admissible directed subset and latent completion. The notation therefore permits feedback without asserting it and permits common-cause association without manufacturing a causal edge.

### 3.4 What “any possible function” means operationally

ECMS is not restricted to linear or monotone relationships. Each active directed component e has a declared admissible function family F_e. F_e may contain positive or negative monotone functions, Hill-like saturation, thresholds, piecewise functions, biphasic or sign-reversing responses, hysteresis, delayed responses, cumulative-exposure functions, persistent reservoirs, recovery functions or other mathematically executable forms. A world selects f_e^(h)∈F_e. The library may be extended whenever the scientific question or evidence requires it.

For reproducibility, “any possible function” cannot mean the unbounded set of all mathematical functions. The method therefore requires an explicit, computationally enumerable grammar. The grammar should be deliberately broad, and evidence should narrow it. If a conclusion changes materially when reasonable alternative function families are added, that dependence is part of the reported uncertainty rather than something to hide.

### 3.5 A complete mechanistic world

For mathematical clarity, one complete world is written

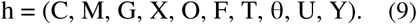

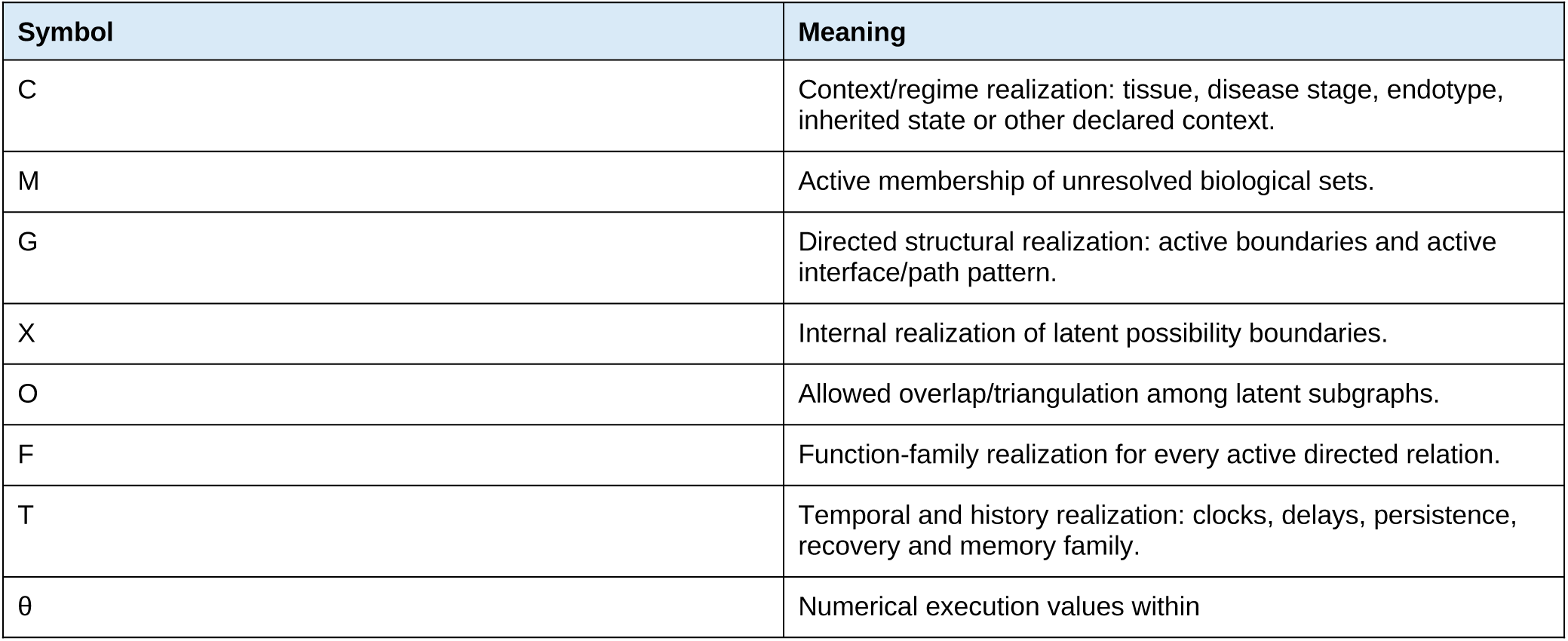
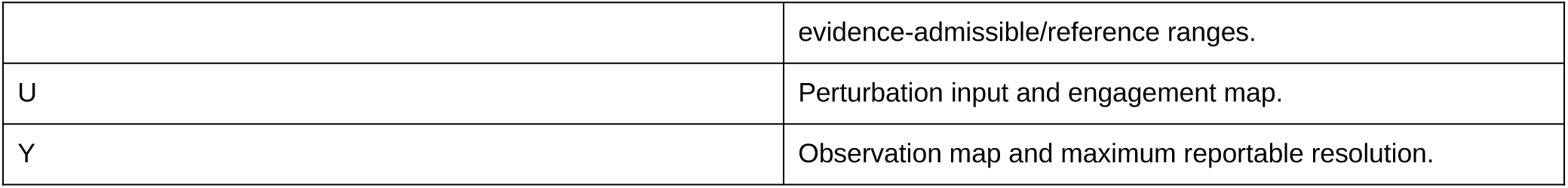

The essential point is that a world contains structural choices as well as numbers. ECMS is therefore not parameter-only Monte Carlo over a fixed graph.

## 4. Step-by-step methodology: Phase I - define the evidence state

### Step 1. Define the scientific question, context of use and version boundary

Before evidence is compiled, specify what the system will be used to ask. Define the disease or biological context, the intervention types of interest, the latent outputs required for the decision problem, the maximum intended observation resolution, and whether the build is prospective or historical. If historical validation is planned, define the knowledge-time cutoff before evidence review. The cutoff applies not only to papers but to concepts, ontology bridges and any external reasoning that can change H(E).

This step prevents “universal model” language from silently widening the scope after construction. A model may be deliberately narrow and still scientifically useful if its context of use is explicit.

### Step 2. Construct the evidence atlas

Collect primary and otherwise admissible evidence with enough provenance to reconstruct every executable statement later. The atlas should preserve source identity, experiment/cohort identity, biological entities, context, intervention/exposure, comparator, timing, endpoint, statistics when recoverable, and a source locator. Reviews may guide search, but executable claims should trace to the underlying evidence or to an explicitly labelled methodological assumption.

### Step 3. Decompose studies into atomic findings without creating pseudoreplication

A publication is not one evidence object. A single experiment may support several statements with different mathematical meanings. Store them as separate atomic findings, but retain a dependency-group identifier whenever rows share participants, samples, experimental material or a tightly linked protocol. Atomicity is about separating information types; it is not a claim of statistical independence.

Example: an experiment may show that intervention P reduces marker A, leaves marker B unchanged and changes disease output only after week 8. These are at least three mathematical statements, but they remain one dependency programme for evidence accumulation unless independence is justified.

### Step 4. Assign information anatomy and dimension-specific transportability

For every atomic finding, ask separately what it identifies about topology, direction, order, bounds, function, magnitude, time/history, context and observation mapping. Then assess transportability separately for each identified dimension. Animal or ex vivo evidence may be valuable for topology or direction while contributing little or no human disease-scale magnitude or kinetics. A human biomarker study may identify an observation ordering without identifying a causal relation. Transportability is therefore a vector of permissions, not one global quality label.

#### Governance note: LLM-assisted semantic compilation and biological intuition

ECMS may use a large language model as an implementation aid for semantic compilation, ontology bridging and biological-intuition candidate generation, but the model is never a scientific evidence source. All source material supplied to the model must obey the active knowledge-time boundary. The model may propose candidate information-anatomy classes, unresolved-set splits, possibility-boundary realizations, function/time/history families or natural-language rationales; every accepted proposal must remain traceable to the supplied evidence and must be adjudicated by a human investigator against the source. LLM assistance must not manufacture numerical magnitudes, target-engagement equivalence, patient-response probabilities or post-cutoff knowledge. Model identity, execution window, task restrictions, accepted frozen rule and human adjudication should be logged. An iterative workflow is permitted: individual prompts are implementation scaffolding rather than scientific evidence, so reproducibility rests on the disclosed source set, governing instructions, accepted-rule log, frozen registries and executable package. Prompt transcripts may be retained or shared for audit when useful, but absence of a public verbatim prompt corpus does not convert LLM output into evidence.

## Step 5. Separate hard boundaries from fallible support

A hard fence is a statement that no retained world is allowed to violate within the declared context. Examples include logical incompatibilities, experimentally established impossibilities at the modelled resolution, or explicit semantic constraints such as “this clinical scale increases when disease activity decreases.” Strong but fallible evidence remains soft. Hard facts should not be represented as q=1 merely to force the sampler; doing so conflates scientific impossibility with high support.

## 5. Step-by-step methodology: Phase II - compile the mechanistic possibility space

### Step 6. Define unresolved biological sets before defining a fixed node ontology

Create an object registry in which every major biological label is initially treated as a set. Define its membership boundary and what kinds of subcomponents could plausibly belong to it. The registry should be broad enough to avoid excluding plausible mechanisms merely because the literature uses inconsistent nomenclature.

### Step 7. Apply the behavioral split criterion

Split a set only when evidence requires behavior that cannot be represented by one unresolved object without losing an executable distinction. Typical triggers are opposite direction, route selectivity, differential perturbation response, different time/history behavior, different function class or context-specific presence. If a context gate is sufficient, prefer a context gate over duplicating the biological object. The burden of proof is on added resolution, because every split removes equivalence and changes the possible-world space.

### Step 8. Create bidirectional possibility boundaries between sets

For every evidence-relevant relationship between unresolved sets, create a boundary A↔X_AB↔B. Define the admissible latent-completion grammar X_AB, the candidate directed interfaces D_AB, and any hard restrictions on those interfaces. Do not assume that an observed influence is direct. Do not assume that the reverse direction is active. Do not prohibit feedback merely because the source reported only one direction. The boundary is a container for all evidence-admissible directed realizations; individual worlds make the directional commitment.

Where a mediator or direct relation is well resolved, include that resolved realization in X_AB. Where mediation is unknown, retain unresolved completions. Where a pure association is all that is supported, include common-cause/observation realizations and do not require a directed A→B path.

### Step 9. Register overlaps and triangulation explicitly

Two possibility boundaries may share hidden biology. If X_AB and X_AC can overlap, record the overlap as an explicit world-level structural option O rather than assuming either complete independence or identity. Transfer only the information dimension justified by the overlap: shared topology does not automatically transfer numerical gain or time. When overlap is proposed by biological intuition rather than a direct source, record that epistemic status separately; see Appendix A.

### Step 10. Encode contradictions as alternative regimes rather than averaged coefficients

Contradictions are information. If studies agree on direction but differ in magnitude, retain the relation and broaden magnitude/function uncertainty or context dependence. If credible evidence supports opposite signs, different selectivity or different sufficiency, review whether a set, context or latent mechanism must split. A negative clinical result with uncertain exposure should constrain the input map before it is allowed to delete the mechanism. A proximal biological effect without downstream clinical improvement should preserve the proximal effect and decouple the downstream claim.

### Step 11. Build the temporal and history grammar

ECMS does not assume one universal clock. Define clock bands and literal evidence boundaries for fast signalling, intermediate recruitment/remodelling, slow structural change or other system-specific scales. If the evidence identifies only that an effect appears by a time or persists beyond a time, encode the inequality rather than fitting a decay curve. History-dependent alternatives may include persistent reservoirs, cumulative exposure, delayed recovery, refractory states, hysteresis or prior-state dependence.

The reduced ECMS state need not be Markovian. For an active directed component e in world h, a general relation contribution can be written

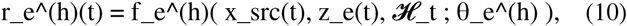

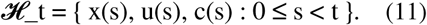

Thus the present response may depend on past states or exposures even when the current perturbation is zero. One executable implementation is a memory kernel

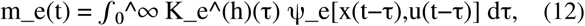

with r_e(t) depending on m_e(t). Another is an augmented hidden reservoir state. These are mathematically different implementations of the same evidence-level statement: history matters. A specific memory mechanism should be selected only if evidence distinguishes it.

### Step 12. Register comparative information as first-class world events

Comparisons should be encoded before independent sampling destroys them. Define pairwise orders, ordinal regimen orders, route-selectivity patterns, temporal orders, crossovers, joint multi-node patterns, non-sufficiency/decoupling and explicit abstention events. A comparison event C_j(h) is evaluated on the complete world under matched context, time and input semantics.

Cross-molecule potency comparisons require comparable proximal engagement. Nominal dose in milligrams is not a common mechanistic intensity scale. Within-molecule dose or regimen order may be constrained without assuming a dose ratio equals an engagement ratio.

### Step 13. Build perturbation-input maps

A regimen is translated into a time-dependent input/engagement map U, not inserted directly as a disease-output effect. For regimen r, let

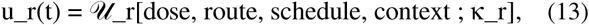

where κ_r contains evidence-supported or residual uncertainty in exposure and target engagement. If proximal engagement is not identified, retain that uncertainty. The same drug can have different input maps by route or regimen. A post-hoc near-class substitution is not permitted merely because two agents share a label.

### Step 14. Build observation maps and output-resolution fences

Latent model state and observed endpoint must be separated. For endpoint k, define an observation object Y_k that states the strongest defensible relationship between latent state x(t) and measured y_k(t):

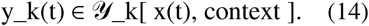

Y_k may be quantitative, bounded, monotone/ordinal, thresholded or merely directional. If only an ordinal relationship is known, the system may compare direction or rank but must not output a calibrated clinical score or patient response probability.

### Step 15. Freeze the possibility-space registries before evidence-allocation compilation

The object registry, boundary/structure registry, function/time grammar, overlap rules, comparison events, input maps, observation maps and hard fences must be frozen before q values are finalized. This separates the scientific question “what worlds are allowed?” from the engineering question “how should supported events be allocated within those worlds?”

## 6. Step-by-step methodology: Phase III - compile evidence allocation and sample worlds

### Step 16. Declare the reference measure μ₀

The hard-constrained possibility space usually remains too large to enumerate. A reference measure μ _o_is therefore required for sampling residual uncertainty. μ _o_should be explicit, simple and sensitivity-tested.

Symmetric optional structural alternatives can use p_0_=0.5; categorical alternatives can use a declared uniform or otherwise justified reference; numerical execution values can use broad bounded ranges; timescales are often sampled on a log scale within evidence-consistent clock bands.

μ _o_is not called a biological prior because ECMS does not require the claim that it represents subjective probability of biological truth. It is the reference against which evidence-induced information is measured.

### Step 17. Convert evidence strength into soft allocation targets q

For each soft event j, define a reference event frequency p_0_,j and compile source support into q_j. The ECMS method requires the adapter to be monotone, dependency-aware, dimension-specific and versioned before final sampling; it does not require one unique numerical formula. A transparent reference adapter is

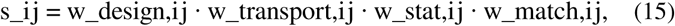

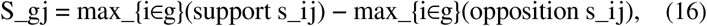

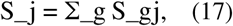

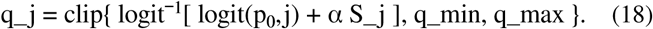

The maxima in Eq. 16 are zero when the corresponding support/opposition set is empty. The grouping g prevents correlated rows from multiplying evidence. Design, transportability, statistics and event match are evaluated for the particular information dimension being allocated. For context-gated evidence, the target is conditional: E_Q[C_j | A_j=1]=q_j, where A_j is the event’s applicability indicator. q is an ensemble-allocation target, not a patient response rate, posterior truth probability or prevalence.

The exact α, caps and within-program aggregation are implementation choices that must be prespecified and sensitivity-tested. Hard fences are not generated by pushing q to 1.

### Step 18. Generate complete candidate worlds hierarchically

Candidate generation is hierarchical because later uncertainties only exist conditional on earlier structural choices. A useful factorization of the reference generator is

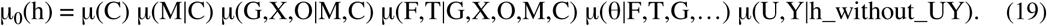

This factorization is conceptual; implementations may regroup factors provided they preserve the same conditional logic. The operational order is: context/regime → active set members → structural/boundary realization → latent completion and overlap → function family → time/history family → numerical execution values → input and observation maps → hard-fence check.

**Figure 3.**
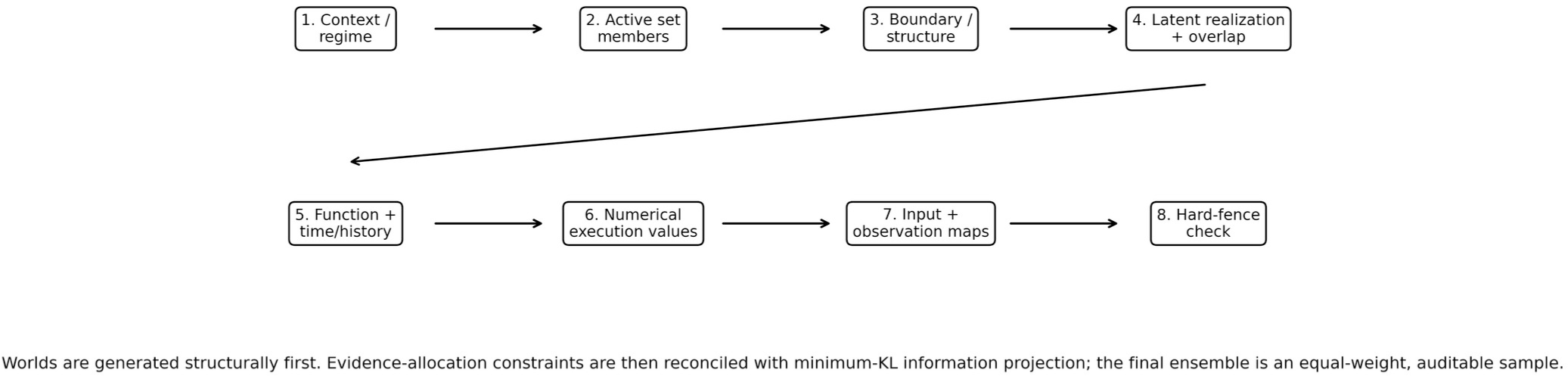
Hierarchical Monte Carlo world generation. The hierarchy prevents a numerical parameter prior from silently deciding whether a biological mechanism exists. “Hierarchical” here describes the generative factorization; it does not imply a Bayesian hierarchical population model.

### Step 19. Use q-guided proposals only to obtain adequate coverage

A neutral generator can make a high-information event extremely rare even when the literature says it should be common. In that case, q-guided proposal, Sequential Monte Carlo (SMC), mutation or other coverage-repair methods may be used to generate enough candidate worlds in both event strata [7]. The proposal mechanism is computational. It must not redefine the scientific possibility space or the final target q.

A rare high-information event should first trigger a scientific check: does the possibility grammar contain the structural/function family required to express the evidence? If not, add the evidence-supported family before using extreme importance weights. Weighting cannot repair a missing mechanism.

### Step 20. Reconcile overlapping soft constraints by minimum-information projection

Let C_j(h) be an event indicator and A_j(h) its applicability indicator. Define the feasible distribution family

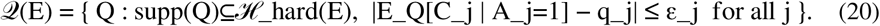

The least-committal evidence allocation is

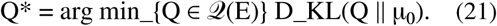

This is an I-projection/minimum-discrimination-information problem [2–4]. For compatible equality constraints, the solution has an exponential-tilt form

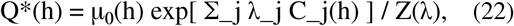

with Lagrange multipliers λ_j chosen to satisfy the registered margins. λ_j are numerical allocation multipliers, not biological parameters. In a finite candidate population, sequential I-projection or iterative proportional fitting can rescale applicable true/false strata repeatedly until all q tolerances are met [4–6]. If the constraints are infeasible, the method must surface the conflict; it must not distort biological parameters until incompatible statements appear to agree.

### Step 21. Audit effective sample size and coverage before equal-weight resampling

For normalized candidate weights w_i, the effective sample size is

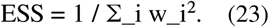

Low ESS signals that the proposal distribution poorly covers the evidence-adjusted target. The correct response is to enrich coverage or use SMC/proposal mutation, not to silently weaken high-information constraints. Every controlling binary event should have support in both true and false strata where both remain admissible.

### Step 22. Create an equal-weight ensemble and choose N empirically

Resample an equal-weight ensemble from the final weighted candidate population for ordinary simulation and retain the weighted pre-resampling population for audit. N is not a defining constant of ECMS. Increase N until event-frequency fidelity, structural coverage, hard-fence compliance, multiseed stability and standardized benchmark-output stability meet prespecified tolerances. The smallest passing N is preferred because it is easier to audit and distribute while preserving the required information.

## 7. Step-by-step methodology: Phase IV - simulate, compare, validate and update

### Step 23. Propagate perturbations through each world without upgrading the observation claim

Simulation proceeds world by world. Let x^(h)(t) be the vector of latent states. A general non-Markovian evolution can be written as a functional differential system

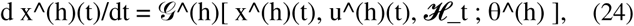

or an equivalent discrete-time/agent/state-transition implementation. The method does not require ordinary differential equations. What matters is that each world uses only its active directed structure, admissible functions, clocks, history rules and input map. Different worlds may use different function families and different temporal mechanisms. This is a feature, not a nuisance: those are unresolved dimensions of the evidence state.

### Step 24. Use a neutral combination rule only when interaction evidence is absent

When multiple fractional perturbations act on the same bounded process and there is no evidence for interaction, a Bliss-style independence reference may be used [12]:

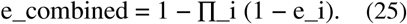

This is a neutral reference, not a claim of synergy. If evidence identifies synergy, antagonism, sequence dependence or pharmacokinetic interaction, encode that information directly. Interaction uncertainty should remain explicit when it matters to conclusions.

### Step 25. Compare regimens on matched mechanistic worlds

Two regimens should be evaluated on the same world h wherever both are executable. Matched-world comparison removes Monte Carlo composition as a source of apparent treatment difference. If R₁ and R₂ are regimens and L_h is a decision-layer loss, define

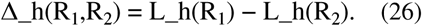

The distribution of Δ_h across matched worlds supports robust ordering, uncertainty and discordant-world analysis. If proximal engagement is not comparable, abstain from a mechanistic potency rank rather than forcing one.

### Step 26. Keep the decision objective downstream of evidence

A researcher may specify a desired target profile over latent outputs. For node j, let d_j denote desired direction, m_j desired normalized magnitude and w_j importance. A generic per-world objective is

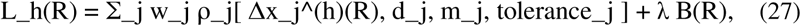

where ρ_j is a prespecified loss and B(R) is an optional burden/complexity term. The ensemble-level objective can use a median, upper quantile, expected loss under the reference allocation or another declared robust functional. Changing d_j, m_j or w_j may change the preferred perturbation, but it must not change H(E), q, world membership or the evidence-derived biology. Decision-layer ranks are objective-conditioned perturbation priorities, not intrinsic molecular potency or clinical treatment recommendations.

### Step 27. Freeze the system before outcome-blind validation

A validation release must freeze the evidence state, possibility-space registries, q compiler, reference measure, software, seeds, finite ensemble and cryptographic hashes before held-out outcomes are used. Test interventions are mapped to frozen input semantics and test classes without consulting downstream outcomes. If the input, history or observation is not identifiable, the correct prediction is abstention or a lower-resolution statement.

### Step 28. Score validation at the information level that was frozen

A sign prediction is a low-information check. More informative tests include ordinal regimen order, temporal maturation, withdrawal persistence, route selectivity, joint multi-node structure, crossover, decoupling and correct abstention. A latent trajectory should not be numerically compared with a clinical endpoint unless an observation map licenses calibration. Validation must therefore score the registered information property, not the most convenient downstream number.

### Step 29. Treat failure as structural information, not as permission to repair the historical model

A mismatch should be classified before changing the model: state-space omission; incorrect direction/function family; missing pharmacologic persistence; context/endotype omission; population heterogeneity; incorrect input mapping; incorrect observation mapping; or study/reporting bias. A genuine failure in a retrospective outcome-blind historical reconstruction remains a failure of the frozen release. New evidence can motivate a new version; it must never be back-projected into the old release and relabelled as validation success.

### Step 30. Update by reconstructing H(E), not by editing sampled worlds

If new evidence e_new becomes available, the scientific learning operation is

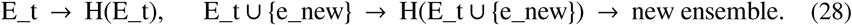

The old frozen ensemble remains an immutable audit snapshot. Learning changes the possibility-space boundaries, constraints or allocation and then resamples a new ensemble. It does not “correct” selected members of an old ensemble in place.

## 8. Interpretation and reporting rules

The following reporting rules are part of the method, not stylistic preferences:

- An ECMS world is an epistemic mechanistic hypothesis. It is not a patient unless a separate patient-population model has been constructed.
- An ensemble frequency is conditional on the declared possibility space, reference measure and evidence-allocation rules. It is not automatically a posterior biological probability.
- q is a target marginal frequency for an evidence event; it is not a clinical response rate or prevalence.
- A↔X↔B is a bidirectional possibility boundary. It does not assert universal feedback. The directed realization is chosen at world level under evidence constraints.
- X is a family of latent subgraph completions, not one hidden biological node.
- Near-zero numerical gain is not equivalent to structural absence; relation presence and numerical intensity should be distinct variables.
- Nominal drug dose is not normalized target engagement across molecules.
- A biomarker association does not become a causal edge unless causal/perturbational evidence identifies that direction.
- Latent output can be reported only at the resolution licensed by the observation map.
- Abstention is a scientific output when the requested comparison is not identifiable.
- Robust conclusions should be checked under reasonable alternate reference measures, q-adapter strengths and function families when those choices could change the conclusion.

## 9. Method invariants, implementation choices and reproducibility contract

Retrospective transparency is an audit device, not a mathematical proof of investigator innocence. Historical source restrictions, LLM decision logging, independent source re-checking, outcome-blind freezes and immutable checksums reduce and expose routes of leakage; they cannot recreate the epistemic state of a genuinely prospective preregistration.

A reader-facing methodology must distinguish principles that define ECMS from engineering choices that can legitimately change across implementations.

| Status | Statement |
| --- | --- |
| Method invariant | Minimum commitment: each finding constrains only the mathematical information it identifies. |
| Method invariant | Biological labels are unresolved sets until a behavioral split is justified. |
| Method invariant | Unresolved relationships are represented as bidirectional possibility boundaries $A \leftrightarrow X \leftrightarrow B$; individual worlds instantiate directed realizations. |
| Method invariant | Association, causal direction, observation mapping and context are distinct objects. |
| Method invariant | Hard fences and soft q allocation are different semantics. |
| Method invariant | World generation is structural/hierarchical before numerical uncertainty. |
| Method invariant | Overlapping soft constraints are reconciled with a least-committal/minimum-information allocation relative to a declared reference measure. |
| Method invariant | Finite ensemble size is chosen by information-preservation/convergence, not convention. |
| Method invariant | Decision preferences remain downstream of evidence. |
| Method invariant | Historical releases are immutable; new evidence creates a new evidence state and ensemble. |
| Implementation choice | Exact function library and numerical ranges, provided they are explicit, evidence-admissible and sensitivity-tested. |
| Implementation choice | Exact evidence-strength adapter, $\alpha$ and q caps, provided they are monotone, dependency-aware and prospectively frozen. |
| Implementation choice | Candidate generator: direct Monte Carlo, SMC, mutation, quasi-Monte Carlo or hybrid methods, provided the target constraints and audits are preserved. |
| Implementation choice | Numerical solver and bounded-state aggregation mechanics. |
| Implementation choice | Decision-layer loss and robust aggregation, provided they do not alter evidence construction. |
| Implementation choice | Exact N and tolerance values, provided selection rules are prespecified and convergence is demonstrated. |

### 9.1 Minimum reproducibility artifacts

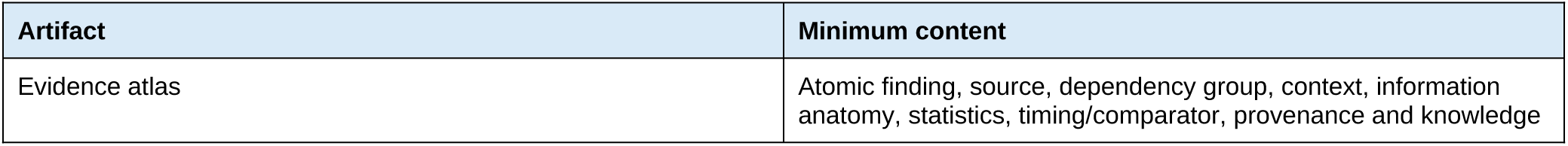
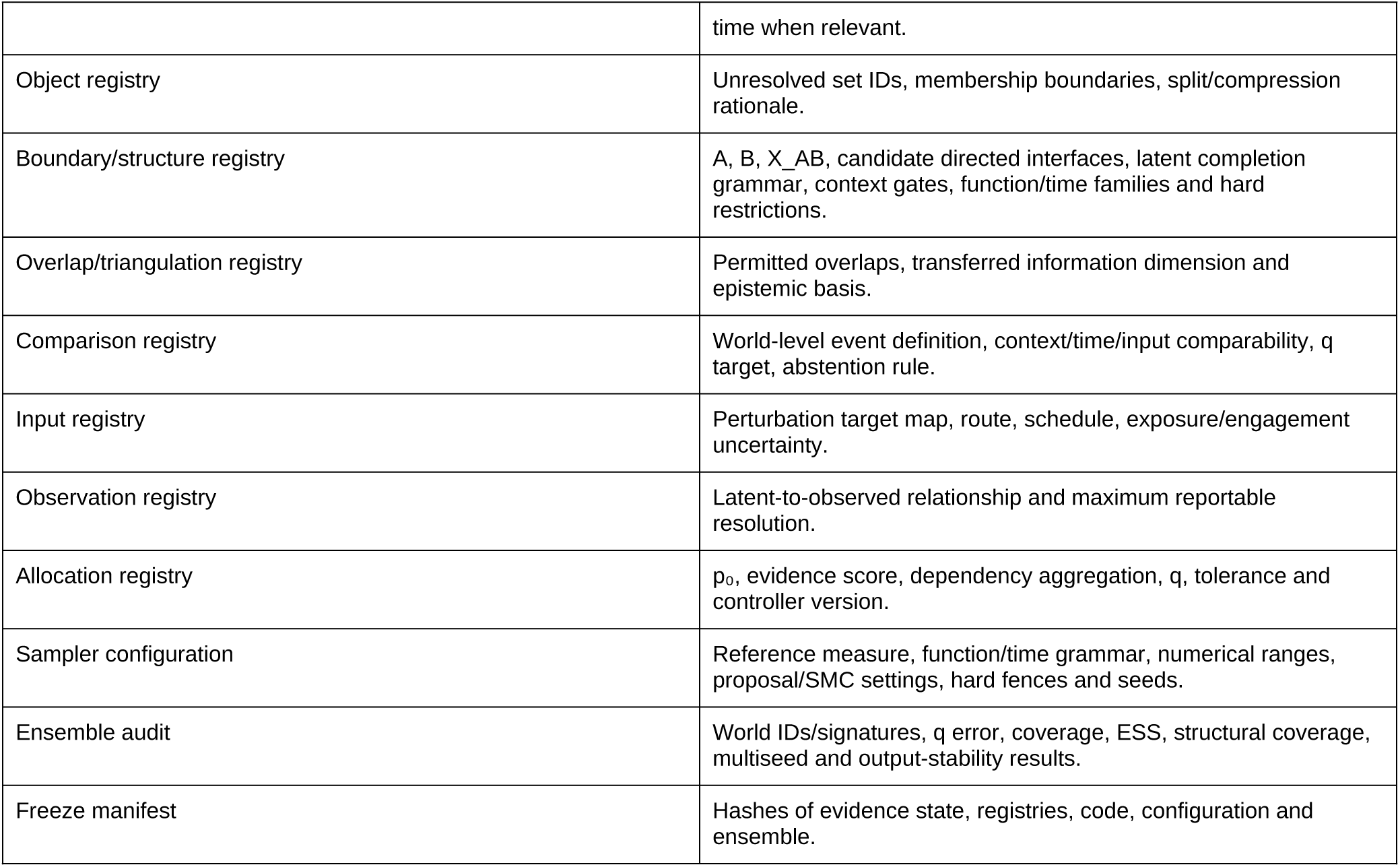

## Appendix A. Glossary of ECMS terms

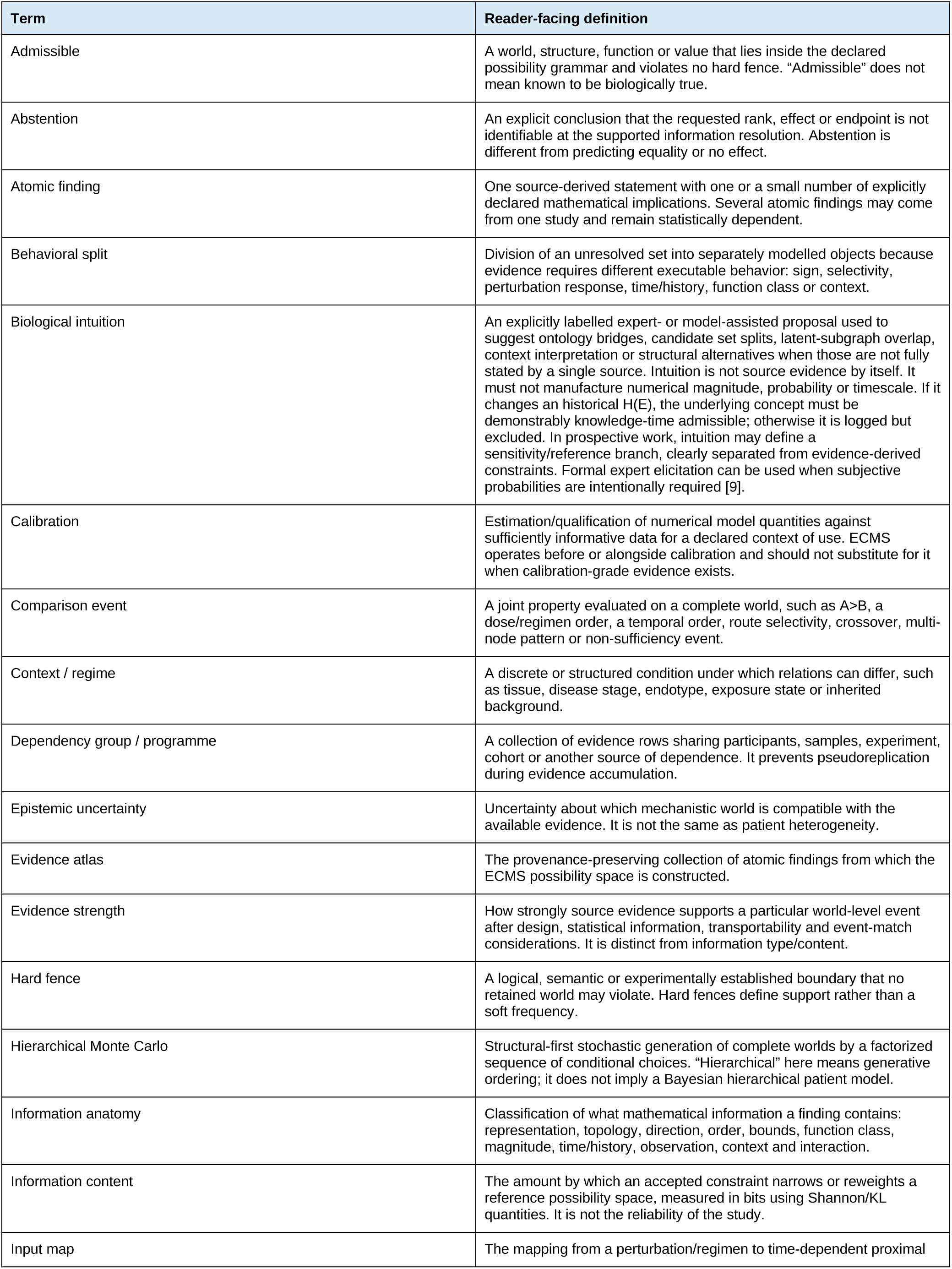
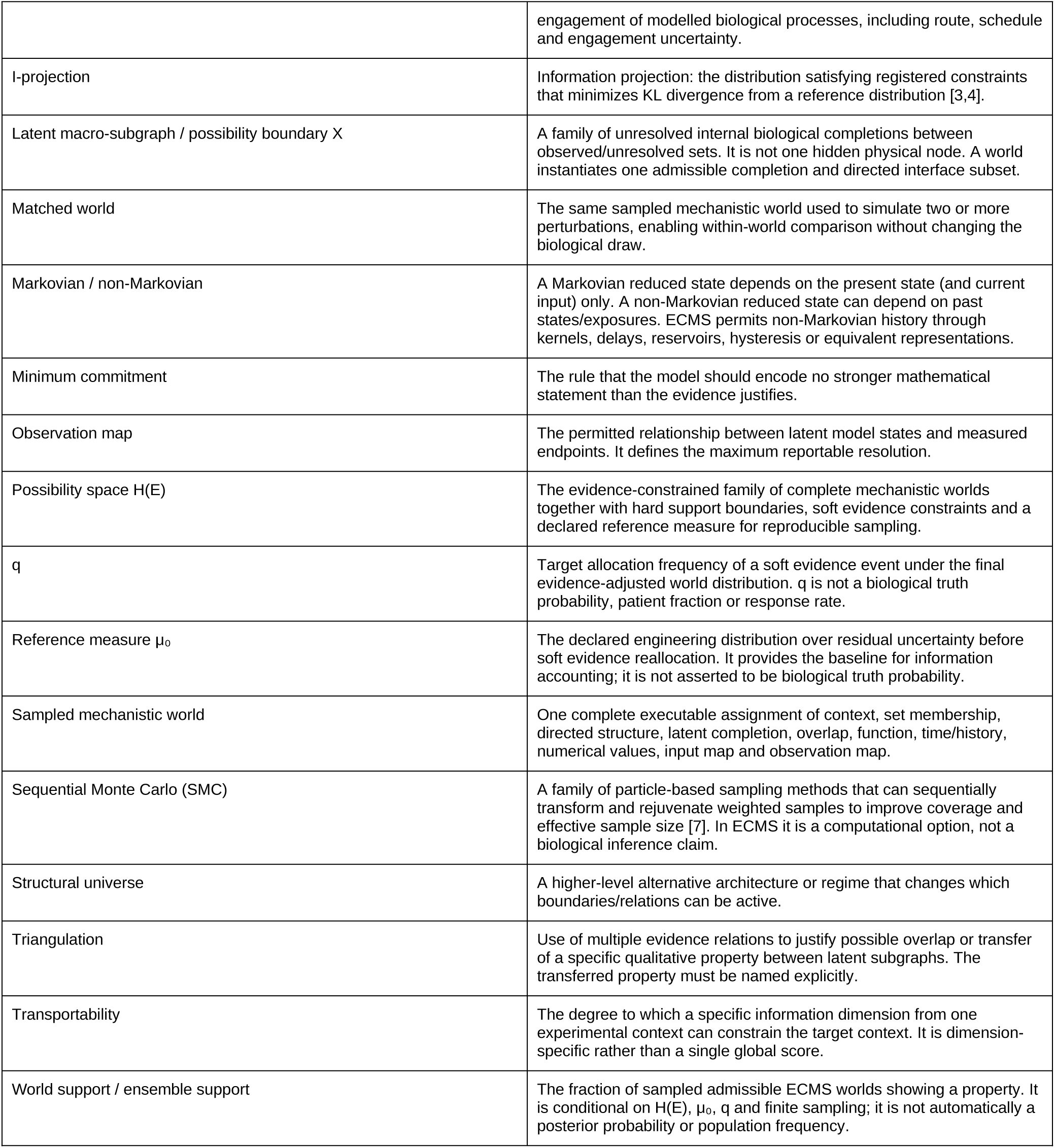

## Appendix B. Abbreviations and symbols

| Abbreviation / symbol | Meaning |
| --- | --- |
| ECMS | Evidence-Constrained Mechanistic Synthesis |
| QSP | Quantitative Systems Pharmacology |
| KL | Kullback-Leibler |
| I-projection | Information projection / minimum-KL projection |
| IPF | Iterative Proportional Fitting |
| SMC | Sequential Monte Carlo |
| ESS | Effective Sample Size |
| LLM | Large Language Model |
| ODE | Ordinary Differential Equation |
| PK/PD | Pharmacokinetics / pharmacodynamics |
| DDI | Drug-drug interaction |
| H(E) | Evidence-constrained mechanistic possibility space |
| $\mu_0$ | Reference measure on admissible worlds |
| $Q^*$ | Least-committal evidence-adjusted world distribution |
| $p_0$ | Reference frequency of a soft event |
| $q$ | Target evidence-allocation frequency |
| $X_{AB}$ | Latent possibility boundary / family of admissible macro-subgraph completions between sets A and B |
| $\mathcal{H}_t$ | Relevant history before time t |
| U | Perturbation/input mapping component of a world |
| Y | Observation-mapping component of a world |

## Appendix C. Toy worked example of a bidirectional possibility boundary

Suppose a literature corpus refers to two coarse biological labels A={a₁,a₂,a₃} and B={b₁,b₂}. The mediator is unresolved. ECMS first declares the boundary A↔X_AB↔B and a broad reference grammar. Four illustrative worlds are shown below; real systems can contain many more combinations.

| World | Active A | Active B | Directed realization | Function family | Time/history | Interpretation |
| --- | --- | --- | --- | --- | --- | --- |
| W1 | $\{a_1, a_2\}$ | $\{b_1\}$ | $A \rightarrow X \rightarrow B$ | positive saturating | short lag | A can drive B. |
| W2 | $\{a_2, a_3\}$ | $\{b_1, b_2\}$ | $B \rightarrow X \rightarrow A$ | negative threshold | intermediate | Reverse influence only. |
| W3 | $\{a_1, a_3\}$ | $\{b_2\}$ | $X \rightarrow A$ and $X \rightarrow B$ | shared latent driver | slow | Association/<br>common cause<br>without $A \rightarrow B$. |
| W4 | $\{a_1, a_2, a_3\}$ | $\{b_1, b_2\}$ | $A \leftrightarrow X \leftrightarrow B$ | forward positive;<br>reverse negative | persistent /<br>hysteretic | Feedback with<br>memory. |

Now suppose evidence provides three statements: (i) perturbing A changes B in the positive direction in a declared context; (ii) A-response precedes B-response; and (iii) an observational dataset shows A and B can covary when the perturbation is absent. Statement (i) increases allocation to worlds with a directed A→…→B path but does not delete reverse or common-cause possibilities unless the evidence is hard. Statement (ii) further constrains temporal families of the forward path. Statement (iii) can preserve or increase shared-latent realizations without being interpreted as a new B→A causal path. If later perturbation evidence establishes feedback, the possibility boundary already has the representational capacity to express it; a new version of H(E) can then reallocate/reshape the world space.

The toy example illustrates why the bidirectional boundary is a possibility envelope rather than a causal conclusion. Direction is a property of the realized world and its evidence events.

## Appendix D. Anti-misinterpretation guide

| Expression | Common but incorrect reading | ECMS reading |
| --- | --- | --- |
| $q=0.80$ | Incorrect: "the mechanism is 80% likely to be true." | Correct: the compiled evidence targets the event in 80% of applicable sampled worlds relative to the declared reference measure. |
| 75% of worlds improve | Incorrect: "75% of patients will respond." | Correct: 75% of sampled epistemic worlds show the property; patient heterogeneity requires a separate population model. |
| $A \leftrightarrow X \leftrightarrow B$ | Incorrect: "A and B necessarily cause one another." | Correct: both directional interfaces are available inside the possibility boundary; each world activates only an admissible directed subset. |
| X | Incorrect: "X is one unknown mediator." | Correct: X is a family of possible latent completions, potentially with different internal structure, overlap, functions and clocks. |
| Near-zero gain | Incorrect: "the edge is absent." | Correct: structural absence and weak numerical intensity are different states unless explicitly collapsed by design. |
| 4 mg vs 50 mg | Incorrect: "4 mg is weaker/stronger mechanistically than 50 mg." | Correct: nominal dose across molecules is not a common target-engagement scale. |
| Biomarker correlates with disease | Incorrect: "biomarker causes disease." | Correct: represent an observation/common-cause relation unless causal perturbation evidence identifies direction. |
| Two perturbations together | Incorrect: "the combination is synergistic." | Correct: use a neutral independence reference if no interaction evidence exists; synergy/antagonism requires evidence. |
| Model cannot rank two regimens | Incorrect: "the regimens are equal." | Correct: the requested order is not identifiable under the current input/observation/evidence state; abstain. |
| Later evidence contradicts freeze | Incorrect: "repair the old world ensemble and rerun validation." | Correct: retain the mismatch, build H(E<sub>new</sub>) as a new version, and preserve the old freeze for audit. |

## Appendix E. Mathematical notes on information projection and convergence

### E1. Conditional evidence margins

Context-specific evidence should not alter worlds outside its scope. If A_j(h) indicates applicability and C_j(h) indicates the evidence event, the target is

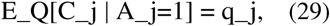

not E_Q[C_j]=q_j over the entire ensemble. The frequency of the context A_j under μ_o_/Q* is itself a reference allocation unless patient prevalence has been separately identified.

### E2. Finite-candidate sequential I-projection

Given candidate weights w_i and an event j, let T be the applicable candidates satisfying C_j=1 and F the applicable candidates satisfying C_j=0. A single exact projection rescales weights within T and F so that their conditional mass becomes q_j and 1−q_j while leaving ratios within each stratum unchanged. Cycling over all events is iterative proportional fitting / sequential I-projection. Convergence is checked numerically; conflicting constraints are reported rather than hidden [4–6].

### E3. Why hierarchical generation is necessary

Flat parameter sampling is inappropriate when some numerical parameters exist only if a structure is active. If relation presence R is uncertain and gain g is drawn first, the numerical prior on g can inadvertently decide what “absence” means. ECMS samples R first, then draws g conditional on R=1. This preserves the difference between structural uncertainty and numerical uncertainty.

### E4. Convergence diagnostics

At minimum, report candidate coverage, weighted ESS, maximum q error, hard-fence violations, structural-signature coverage, multiseed reproducibility and stability of prespecified benchmark summaries as N increases. Reference-measure and evidence-adapter sensitivity should be included when key conclusions depend on them. There is no universal ECMS ensemble size.

### E5. Falsifiability boundary

The latent possibility boundary must not become an unlimited escape hatch. X_AB, F_e and the temporal/history grammar are declared before validation, and hard fences constrain them. A later observation is a genuine failure when reproducing it would require a structure, function, time/history behavior, input map or observation map that was outside the frozen admissible space. The correct scientific response is to record the failure and determine which coordinate should change in the next evidence state.

## Methodological status

This document defines the reader-facing ECMS methodology. Disease-specific evidence, fitted/normalized values, frozen ensembles and validation outcomes are implementation artifacts and should be cited separately in disease applications, not used as foundational references for the method itself.

## Supplementary Information

Evidence-constrained mechanistic synthesis for pre-calibration perturbation inference Supplementary Note 1 is supplied as a separate authoritative Supplementary Methodology document. This file contains Supplementary Notes 2-13 and Tables S1-S6. Tables S7-S19 are in the accompanying Supplementary Tables workbook.

### Supplementary Note 2 | Historical evidence reconstruction and knowledge time

The historical evidence state was defined independently for alopecia areata (AA) and chronic spontaneous urticaria (CSU) using information available on or before 31 December 2023. The knowledge-time rule applied to any information capable of changing H(E), including papers, registry results, conference disclosures, ontology bridges, biological intuition and external reasoning. Journal publication year was therefore not sufficient to establish that a result was unseen.

LLM-assisted semantic compilation in this study. From 1 to 20 August 2026, the historical packages were constructed iteratively with OpenAI GPT 6-sol thinking models operated at high reasoning effort. For construction tasks, the models were given pre-cutoff evidence material and ECMS information-anatomy/governance instructions and were used to propose source-grounded semantic translations, ontology bridges and biological-intuition candidates. They were not treated as evidence, were not authorized to use held-out post-cutoff outcomes for construction decisions, and did not make final scientific acceptance decisions. Investigators manually checked proposals against cited sources, accepted/revised/rejected the proposals and manually verified both the frozen packages and the manuscript-facing intuition log. The workbook sheet LLM_Biological_Intuition_Log (Supplementary Table S18) records 84 accepted q-event rules with linked historical evidence identifiers, model family, execution window, prohibitions and human adjudication. Individual iterative prompts are not disclosed as part of the public scientific evidence package; partial prompt traces can be shared with reviewers if specifically useful. This audit improves traceability and reduces leakage risk but does not prove that retrospective investigator bias was impossible.

Search and adjudication proceeded in layers: broad disease/mechanism search, focused perturbation and observation-map searches, source-level dependency grouping, atomic finding extraction, contradiction/context coding and calibration-readiness review. Multiple readouts from one experimental programme remained dependent unless a finer independence structure was justified. The complete locked evidence atlases, search logs and dependency registries are supplied as Supplementary Data rather than reproduced in this narrative.

Post-cutoff candidate studies were screened using the same knowledge-time rule. Studies whose relevant results had been publicly disclosed before the cutoff were retained only as compatibility/training-overlap records and excluded from unseen validation. Representative knowledge-time decisions are shown in Supplementary Table S2.

**Supplementary Table S1.**
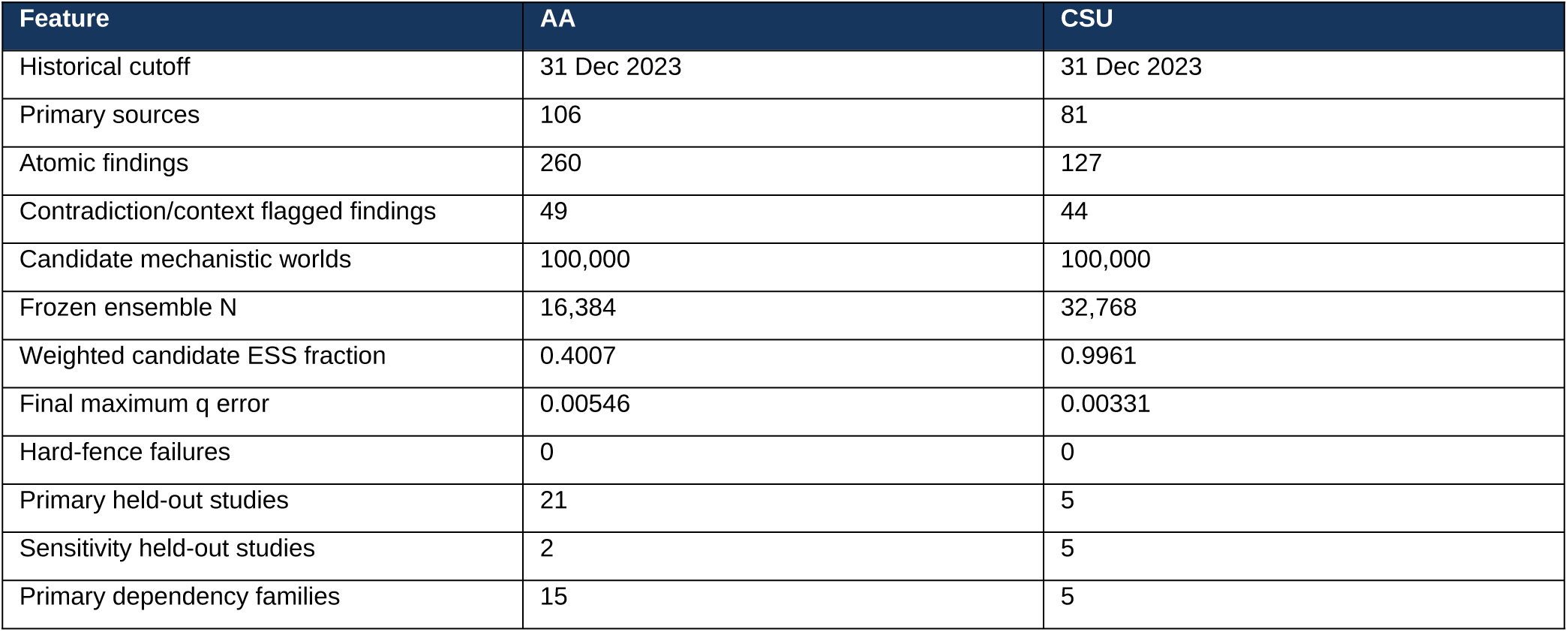
Historical construction and freeze summary.

| Feature | AA | CSU |
| --- | --- | --- |
| Historical cutoff | 31 Dec 2023 | 31 Dec 2023 |
| Primary sources | 106 | 81 |
| Atomic findings | 260 | 127 |
| Contradiction/context flagged findings | 49 | 44 |
| Candidate mechanistic worlds | 100,000 | 100,000 |
| Frozen ensemble N | 16,384 | 32,768 |
| Weighted candidate ESS fraction | 0.4007 | 0.9961 |
| Final maximum q error | 0.00546 | 0.00331 |
| Hard-fence failures | 0 | 0 |
| Primary held-out studies | 21 | 5 |
| Sensitivity held-out studies | 2 | 5 |
| Primary dependency families | 15 | 5 |

**Supplementary Table S2.** Representative knowledge-time adjudications.

| Record | Earliest public availability | Disposition |
| --- | --- | --- |
| CSU remibrutinib REMIX programme | 9 Aug 2023 sponsor topline | Pre-cutoff compatibility only |
| CSU ligelizumab PEARL programme | 23 Nov 2023 e-publication | Pre-cutoff compatibility only |
| CSU ARROYO | 9 Nov 2023 registry results | Pre-cutoff compatibility only |
| AA THRIVE-AA1 | 2022 public conference disclosure | Training overlap / compatibility only |
| AA ALLEGRO-LT month 24 | 2022 public conference abstract | Training overlap / compatibility only |

### Supplementary Note 3 | Disease-specific implementation of the general ECMS grammar

The general methodology in Supplementary Note 1 was instantiated separately for AA and CSU. Disease-specific labels are compressed biological coordinates, not claims that each label is an indivisible molecular entity. Each coordinate can represent an unresolved set whose members remain compressed unless evidence demonstrates behaviorally different executable consequences.

AA used a follicle-centred immune-to-output grammar with historical drivers and dynamic coordinates representing immune-privilege/visibility, innate/type-I inflammatory activity, JAK-sensitive cytokine signalling, recruitment, regulatory and cytotoxic effector functions, and follicular integrity/hair output. CSU used autoallergic, type-IIb autoimmune and other activating contexts together with dynamic Fc-epsilon-RI competence, proximal signalling, complement, mast-cell competence/effector activity, eosinophilic/cytokine support, coagulation/contact-system activity and clinical wheal/itch output. Disease-specific object, boundary, relation, input and observation registries are supplied in Supplementary Data.

The main manuscript uses biological descriptions rather than implementation variable codes. Stable IDs are retained only for reproducibility and in the supplementary tables.

### Supplementary Note 4 | Evidence strength, dependency aggregation, q and information projection

Information type and evidence strength were kept distinct. A finding first determines what mathematical object may be constrained. Evidence reliability and independent replication then determine how strongly the corresponding event is allocated across candidate worlds. Dependent findings are aggregated within dependency programmes before independent programmes are combined. This prevents one source with many readouts from acting as many independent replications.

Soft evidence was compiled as target event allocations q relative to a declared reference measure. q is a construction frequency, not posterior biological truth. Candidate-world weights were adjusted by sequential minimum-information projection until the registered conditional margins were approached within tolerance. Infeasible or conflicting margins would be reported; neither historical system required silent constraint deletion. Complete q-event definitions, evidence identifiers, dependency programmes and realized frequencies are available in Supplementary Table S9 and the frozen audit files.

Targeted adapter sensitivity used the prespecified evidence-strength slope values alpha=1.2, 1.6 and 2.0 on the same scientific grammar. The favoured side of registered controllers did not reverse across this band; only allocation strength changed. This is an allocation-target sensitivity analysis, not a confidence interval and not a replacement ensemble unless the whole reallocation/simulation is rerun.

### Supplementary Note 5 | World generation, function/time/history coverage and ensemble adequacy

Candidate worlds were generated hierarchically: context and unresolved-set membership -> possibility-boundary realization -> latent completion and overlap -> function/sign -> time/history family -> numerical execution values -> perturbation input map -> observation map -> hard-fence audit. This preserves the distinction between structural absence and weak numerical effect.

The finite grammar admitted multiple nonlinear and temporal behaviours. The frozen AA active relation instances included linear, power, Hill-like, threshold, piecewise, weak-linear and gating forms; CSU additionally represented delay and persistent forms. Each disease also retained exponential, ramp and step temporal families and four whole-world history families. These frequencies are a coverage census of the frozen epistemic ensemble; they are not biological prevalence.

Adaptive N selection used evidence-allocation fidelity, hard-fence compliance, multiseed reproducibility and stability of prespecified benchmark summaries on a common standardized scale. In CSU, N=16,384 failed the three-seed stability criterion (maximum standardized summary difference 0.01893); N=32,768 was the smallest passing tier (0.00394). AA froze at N=16,384 after its own prospective audit.

### Supplementary Note 6 | Perturbation inputs, observation maps and abstention

Input maps were molecule/regimen specific. Nominal doses from different molecules were never interpreted as a common normalized engagement scale. Route-sensitive inputs, including topical versus systemic JAK perturbation, were kept separate when tissue exposure changed the biological interpretation. Mechanistic inputs such as MRGPRX2 blockade remained mechanistic unless a clinical bridge was independently supported.

Observation maps specify the resolution at which a latent variable can be compared with a measured outcome. AA hair-output trajectories were evaluated qualitatively/ordinally against SALT or local hair response; CSU latent activity was evaluated qualitatively against UAS7 and inversely against UCT. No calibrated patient-response probability was created. When intervention, history, comparison or observation resolution was not identified, the registered output was abstention or a lower-resolution statement.

### Supplementary Note 7 | Validation firewall and concordance grading

Atomic reporting is separated into empirical answers, falsification, unassessable committed propositions and abstention. Across 41 registered atomic properties, ECMS made 34 executable commitments: 29 were empirically concordant, one was a clear mismatch, and four could not be fully assessed because the public held-out record lacked required intermediate values or an isolated intervention component. Seven additional properties were prespecified abstentions and are not counted as empirical successes. The Reporting_Summary sheet in the master workbook preserves these counts.

For each post-cutoff test, intervention identity, route, regimen, evaluable time, observation map and information class were frozen before held-out outcome values were entered. Outcome-blind prediction artifacts were checksummed before scoring. The validation unit is the registered information property, not the most convenient clinical number.

**Supplementary Table S3.** Concordance grades.

| Grade | Operational meaning | Required reason |
| --- | --- | --- |
| Concordant | All assessable registered information properties are supported. | Information class and outcome-source location. |
| Concordant with abstention boundary | Executable property is supported and a prespecified causal/rank/clinical comparison is correctly withheld. | State what was supported and what was intentionally not claimed. |
| Partially concordant | Principal property is compatible but a registered component is contradicted or cannot be verified. | Classify as true mismatch, reporting insufficiency, input non-identifiability or observation non-identifiability. |
| Discordant | A registered frozen information property is clearly contradicted. | State the exact proposition that failed; do not repair or average away. |

Study-level labels summarize the registered atomic components. They do not override a discordant atomic proposition: AA-PUB-099 remains study-level partially concordant, while its week-36-to-52 monotonicity component is atomically discordant. Complete study and atomic tables are Supplementary Tables S7 and S8.

### Supplementary Note 8 | Complete AA held-out validation

The AA confirmatory layer comprised 21 primary studies across 15 dependency families; two additional studies were sensitivity analyses. Fourteen primary studies were concordant, four concordant with a prespecified abstention boundary and three partially concordant. No primary study was labelled fully discordant, but AA-PUB-099 contains the explicit atomically discordant week-36-to-52 temporal component.

The most informative AA categories were withdrawal/history, multi-time maturation, local-route transfer and correct non-identifiability. Baricitinib withdrawal (AA-PUB-003) is a cross-stream lineage: same-drug active-response evidence existed before cutoff, but the baricitinib-withdrawal result did not. The frozen history/persistence family was enabled by independent pre-cutoff JAK withdrawal and treatment-history evidence. The full study-level record is Supplementary Table S7; each atomic property is Supplementary Table S8; its evidence ancestry is Supplementary Table S9.

Several low-information active-treatment tests are direct historical analogues or near transfers and should be interpreted as compatibility rather than headline validation. This distinction is encoded explicitly in Supplementary Table S9.

### Supplementary Note 9 | Complete CSU held-out validation

The CSU primary layer comprised five omalizumab 300-mg studies, each in a separate dependency family. Three were concordant and two were partially concordant because the public reports did not expose all intermediate timepoint values required for the registered temporal order. The narrowness of this primary layer is a limitation, not a hidden feature.

Five sensitivity tests supplied broader structure: one omalizumab biosimilar active-ingredient sensitivity, a rapid cyclosporine timing test, two independent MRGPRX2 route-selectivity tests with clinical-extrapolation abstention, and one UCT observation-map recovery. Complete records are Supplementary Tables S7-S9.

### Supplementary Note 10 | Evidence-to-prediction lineage and duplication-risk audit

Each atomic held-out property was backtraced through the frozen construction. The audit records the validation query, input and observation maps, minimum ECMS path, function/time/history property, q events, hard fences, pre-cutoff atomic evidence identifiers, dependency programmes and source identities. It also records whether the same molecule, regimen, endpoint, time/property and exact study/trial programme were present before cutoff. Classification is descriptive and deliberately avoids a numerical novelty score.

**Supplementary Table S4.**
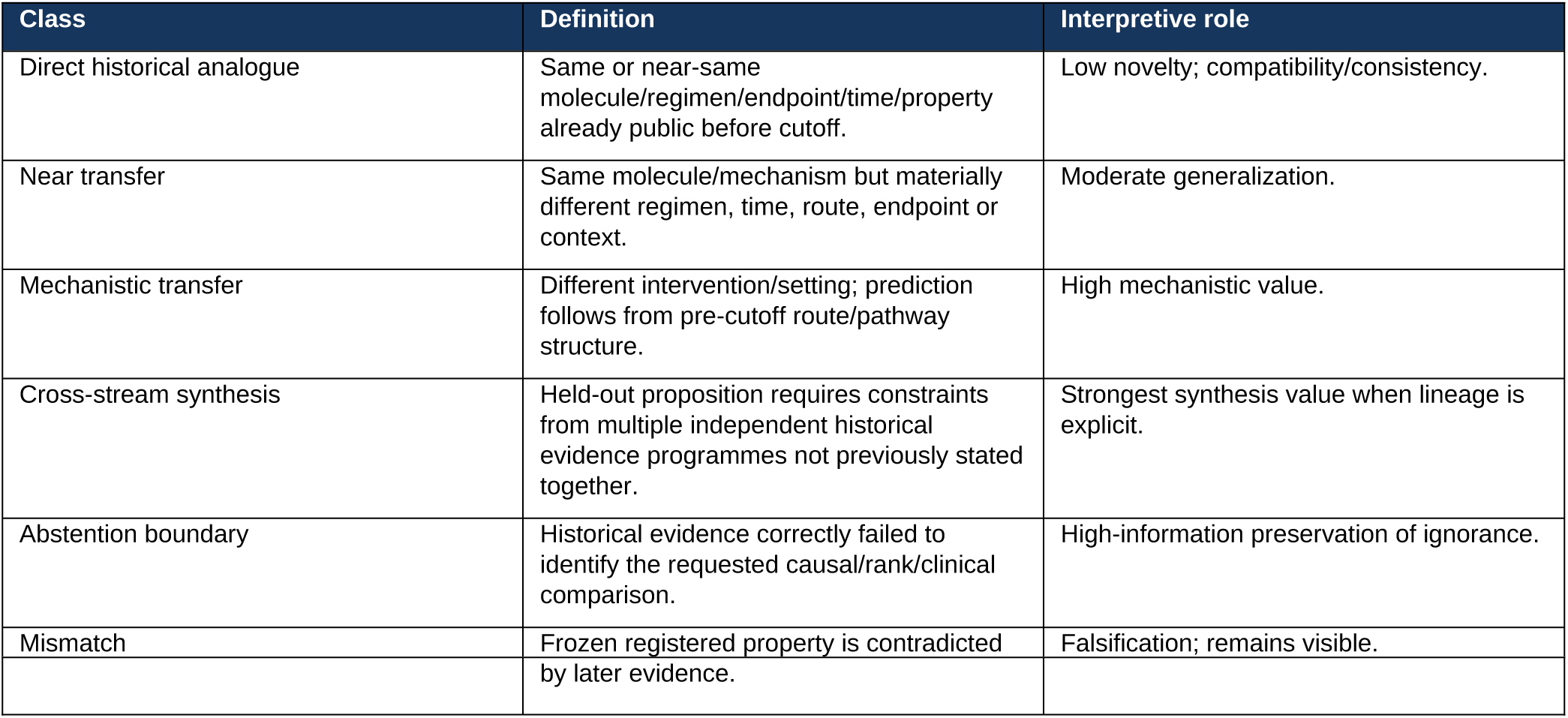
Lineage classes.

Forty-one atomic properties were audited: 9 direct historical analogues, 16 near transfers, 4 mechanistic transfers, 4 cross-stream syntheses, 7 abstention boundaries and 1 mismatch. AA baricitinib withdrawal is cross-stream synthesis; CSU MRGPRX2 antagonism is mechanistic transfer; AA-PUB-099 is the sole explicit mismatch and is notable because the failed late temporal property itself had a direct historical analogue.

Nearest-historical-analogue baseline interpretation. The direct-historical-analogue class is intentionally conservative: it marks properties that a simple same-drug/regimen/endpoint/time matching rule could plausibly reproduce. Near-transfer, mechanistic-transfer and cross-stream classes identify increasing departures from exact historical matching, but this taxonomy is not a formal performance comparison against every possible rule-based, fixed-graph or learned baseline. Such external computational benchmarking remains future work on a predeclared common benchmark.

**Supplementary Table S5.** Representative evidence ancestries.

| Held-out property | Lineage | Pre-cutoff enabling evidence | Interpretation |
| --- | --- | --- | --- |
| AA baricitinib withdrawal | Cross-stream synthesis | Same-drug active BRAVE response + independent JAK withdrawal/history programmes | No pre-cutoff baricitinib-withdrawal result was used. |
| CSU MRGPRX2 antagonism | Mechanistic transfer | Agonist/knockout route separation + mast-cell-to-output bridge | Held-out antagonists test the reverse perturbational direction. |
| CSU omalizumab W12 | Direct historical analogue | Same regimen, clinical endpoint class and W12 dose-order evidence | Compatibility result. |
| AA W36->W52 maturation | Mismatch | Same-regimen historical continued W36->W52 improvement | Later trajectory contradicted the inherited property. |

The complete row-level audit is Supplementary Table S9 in the accompanying workbook. Supplementary Table S10 maps every historical source identity used by those rows.

### Supplementary Note 11 | Robustness and sensitivity analyses

The evidence-to-q adapter is an engineering choice. The present sensitivity analysis varies the prespecified evidence-strength slope alpha while retaining the same source set, dependency aggregation, reference measure, caps and candidate grammar. Stability of the favoured side therefore establishes only a limited robustness claim. Broader sensitivity to alternative reference measures, soft caps and leave-one-dependency-out constructions was not used to redesign the historical packages after outcome reveal and remains an explicit target for future prospective benchmarking.

**Supplementary Table S5a.** Core numerical diagnostics.

| Metric | AA | CSU |
| --- | --- | --- |
| Candidate worlds | 100,000 | 100,000 |
| Frozen N | 16,384 | 32,768 |
| Weighted ESS fraction | 0.4007 | 0.9961 |
| Final max q error | 0.00546 | 0.00331 |
| Final q failures | 0 | 0 |
| Unique frozen candidates | 16,017 | 32,768 |
| Distinct structural signatures | 13,599 | 16,533 |
| Allocation cycles | 24 | 3 |

The AA ESS reflects more concentrated evidence reweighting, but 13,599 structural signatures remained represented after resampling. CSU retained all candidate identities at the frozen tier. Function/time/history family usage and q-adapter sensitivity are reported in Supplementary Tables S12-S15. No frequency in these audits should be read as a patient or biological prevalence.

### Supplementary Note 12 | Decision layer, online research interface and related methods

The decision layer evaluates researcher-declared biological target profiles on the unchanged frozen ensemble. It can compare frozen molecule/regimen inputs and clearly labelled mechanistic inputs. Objective-fit percentages are loss-derived target-profile matches, not potency, efficacy or response probabilities. The public interface placeholder is https://gen-lang-client-0942684525.uc.r.appspot.com/.

ECMS is related to, but scientifically distinct from, several method families. The comparison below is architectural rather than a benchmark; no claim is made that another method ‘cannot’ represent an ECMS feature unless its primary paper establishes that scope.

**Supplementary Table S6.** Neutral architecture comparison.

| Method family / representative papers | Scientific object and evidence regime | Relation / uncertainty emphasis | Relation to ECMS |
| --- | --- | --- | --- |
| Knowledge graphs / graph drug models (PrimeKG; TxGNN) [main refs 11,16] | Large integrated biomedical graphs; learned graph representations can support drug-disease prediction. | Graph relations and learned embeddings; uncertainty follows the model/training formulation. | Complementary: ECMS starts from information-typed literature constraints over multiple complete mechanistic worlds. |
| Perturbation predictors (GEARS; CellOT; scGPT) [12,13,15] | High-dimensional experimental perturbation data and learned response mappings. | Predictive response functions/generalization from measured perturbational datasets. | Complementary when perturbation datasets exist; ECMS addresses heterogeneous literature before commensurate training/calibration data. |
| Causal perturbation / structural-causal models (CINEMA-OT; deep SCMs) [14,17] | Counterfactual or causal treatment-effect objects under explicit identification/model assumptions. | Causal matching or specified/learned structural causal mechanisms. | ECMS does not treat all literature as one causal dataset; causal direction is admitted only at source-supported resolution. |
| Agentic / statement assembly (INDRA) [9,10] | Normalized mechanistic statements assembled from text/databases into executable representations. | Provenance-rich mechanistic knowledge and assembly policies. | Strong precedent for literature-to-executable modelling; ECMS adds information anatomy, possibility-space allocation and first-class abstention. |
| Calibrated mechanistic/QSP [3,4] | Quantitative dynamic models for a defined context of use using mechanistic and PK/PD data. | Calibrated/qualified parameters and explicit pharmacology. | Downstream regime that ECMS is intended to complement, not replace. |

### Supplementary Note 13 | Reproducibility inventory and file identities

The manuscript package contains: (i) this Supplementary Information; (ii) Supplementary Methodology (Note 1); (iii) the master Supplementary Tables workbook; (iv) main and Extended Data figures with source-data CSVs; (v) locked historical evidence atlases; (vi) frozen AA and CSU scientific packages; (vii) validation input/prediction/result reference bundles; and (viii) Supplementary Software 1. Reader-facing aliases are used in the manuscript. Immutable legacy filenames, if present, are retained only in reproducibility manifests with checksum mappings.

The reproducible code/data archive, frozen ensemble arrays and scoring scripts will be linked through the study GitHub repository at https://[ECMS-GITHUB-URL-TO-BE-ADDED] in the final submission. The interactive portal remains a separate research interface at https://gen-lang-client-0942684525.uc.r.appspot.com/.

The accompanying Supplementary Tables workbook uses the following stable numbering: S7 = Concordance_Master; S8 = Atomic_Property_Master; S9 = Lineage_Audit; S10 = Historical_Source_Map; S11 = Lineage_Summary; S12 = q_Alpha_Sensitivity; S13 = Ensemble_Adequacy; S14 = N_Selection; S15 = Function_Time_Census; S16 = Crossref_Map; S17 = Availability_Placeholders; S18 = LLM_Biological_Intuition_Log; S19 = Reporting_Summary.

